# Structural basis for translation inhibition by the glycosylated antimicrobial peptide Drosocin from *Drosophila melanogaster*

**DOI:** 10.1101/2022.12.08.519698

**Authors:** Timm O. Koller, Martino Morici, Max Berger, Haaris A. Safdari, Deepti S. Lele, Bertrand Beckert, Kanwal J. Kaur, Daniel N. Wilson

**Affiliations:** Institute for Biochemistry and Molecular Biology, University of Hamburg, Martin-Luther-King-Platz 6, 20146 Hamburg, Germany; National Institute of Immunology, Aruna Asaf Ali Marg, New Delhi, 110067, India; Dubochet Center for Imaging (DCI) at EPFL, EPFL SB IPHYS DCI, Lausanne, Switzerland

## Abstract

The proline-rich antimicrobial peptide (PrAMP) drosocin is produced by *Drosophila* species to combat bacterial infection. Unlike many PrAMPs, drosocin is O-glycosylated at threonine 11, a post-translation modification that enhances its antimicrobial activity. Here we demonstrate that the O-glycosylation influences not only cellular uptake of the peptide, but also interacts with its intracellular target, the ribosome. Cryo-electron microscopy structures of glycosylated drosocin on the ribosome at 2.1-2.8 Å resolution reveal that the peptide interferes with translation termination by binding within the polypeptide exit tunnel and trapping RF1 on the ribosome, reminiscent of that reported for the PrAMP apidaecin. The glycosylation of drosocin enables multiple interactions with U2609 of the 23S rRNA, leading to conformational changes that break the canonical base-pair with A752. Collectively, our study provides novel molecular insights into the interaction of O-glycosylated drosocin with the ribosome, which provides a structural basis for future development of this class of antimicrobials.

## Introduction

The host defense systems of mammals and higher insects produce a battery of potent antimicrobial peptides (AMPs) in response to bacterial infection. Unlike most AMPs that kill bacteria using a lytic mechanism, proline-rich AMPs (PrAMPs) pass through the bacterial membrane and target intracellular processes, such as protein synthesis (Castle et al., 1999; Graf et al., 2017; Graf and Wilson, 2019; Krizsan et al., 2014; Mardirossian et al., 2014; Scocchi et al., 2011). Two types of PrAMPs have been identified and classified based on their mechanism of action to inhibit protein synthesis, namely, type I PrAMPs that block the accommodation of the aminoacyl-tRNA directly following translation initiation, and type II PrAMPs that do not interfere with initiation and elongation, but prevent dissociation of the release factors RF1 and RF2 during the termination phase (Graf and Wilson, 2019). Structures on the ribosome of a variety of type I PrAMPs from both insect (oncocin, metalnikowin I and pyrrhocoricin) and mammalian (Bac7 and Tur1A) origin have revealed overlapping binding sites that span from the ribosomal exit tunnel to the A-site of the peptidyltransferase center (PTC) (Gagnon et al., 2016; Mardirossian et al., 2018b; Mardirossian et al., 2020; Roy et al., 2015; Seefeldt et al., 2016; Seefeldt et al., 2015). It has been proposed that by occluding the A-site at the PTC on the ribosome, these type I PrAMPs prevent the binding of the aminoacylated CCA-end of the incoming A-site tRNA, and thereby arrest translation (Gagnon et al., 2016; Graf et al., 2017; Graf and Wilson, 2019; Roy et al., 2015; Seefeldt et al., 2016; Seefeldt et al., 2015). Structures on the ribosome with the type II PrAMP Api137, a synthetic derivative of the natural PrAMP apidaecin, have revealed a binding site within the ribosomal exit tunnel that overlaps with type I PrAMPs (Chan et al., 2020; Florin et al., 2017; Graf et al., 2018). However, the binding mode of Api137 is completely different, with a reversed orientation compared to type I PrAMPs, and also Api137 does not encroach so dramatically on the A-site of the PTC. Moreover, Api137 inhibits translation by trapping the termination release factors on the ribosome following peptidyl-tRNA hydrolysis (Florin et al., 2017; Graf et al., 2018).

In addition to the classical membrane-targeting AMPs, such as defensins, cecropins and diptericins, *Drosophila* also produce a PrAMP called drosocin (Bulet et al., 1993; Bulet et al., 1999). Drosocin is 19 amino acids long and, like many PrAMPs, is rich in proline and arginine residues (Bulet et al., 1993) (**Fig. 1a**) and displays excellent activity against Gram-negative bacteria, such as *E. coli* (Bikker et al., 2006; Bulet et al., 1993; Bulet et al., 1999). However, unlike most PrAMPs, drosocin carries an O-glycosylation on residue Thr11, consisting of either the monosaccharide N-acetylgalactosamine (α-D-GalNAc) or a disaccharide comprising galactose linked to an N-acetylgalactosamine (β-Gal(1 →3)-α-D-GalNAc) (**Fig. 1a,b**) (Bulet et al., 1993; Uttenweiler-Joseph et al., 1998). A double glycosylated form of drosocin bearing the monosaccharide on Ser7 as well as Thr11 has also been reported (Rabel et al., 2004). Both the mono- and di-saccharide forms of drosocin appear in Drosophila hemolymph within 6 hours post-infection and increase in concentration (to 40 µM) for up to 24 hours (Uttenweiler-Joseph et al., 1998). While the disaccharide form disappears two weeks after infection, the monosaccharide persists for up to three weeks (Uttenweiler-Joseph et al., 1998). Synthetic drosocin lacking O-glycosylation is less active than the native compounds, suggesting that the post-translational modification is necessary for full activity (Bulet et al., 1993; Bulet et al., 1999; Bulet et al., 1996; Gobbo et al., 2002; Hoffmann et al., 1999). Indeed, many studies have demonstrated that a variety of synthetic drosocin derivatives with varying sugar moieties maintain good antimicrobial activity, generally better than the unmodified form (Ahn et al., 2011a; Ahn et al., 2011b; Gobbo et al., 2002; Lele et al., 2015a; Marcaurelle et al., 1998; Otvos et al., 2000; Rodriguez et al., 1997; Talat et al., 2011). Although NMR and CD experiments suggest that both the modified and unmodified forms of drosocin adopt extended conformations in solution (Bulet et al., 1996; Gobbo et al., 2002; Lele et al., 2015a; McManus et al., 1999; Talat et al., 2011), the presence of the modification has nevertheless been proposed to help drosocin maintain an extended conformation to facilitate binding to its intracellular target (Bulet et al., 1999; Gobbo et al., 2002; McManus et al., 1999). Additionally, glycosylation can also increase solubility, serum stability and broaden the biological activity spectrum (Bulet et al., 1999), however, the exact role of glycosylation for drosocin remains unclear.

**Fig. 1:**
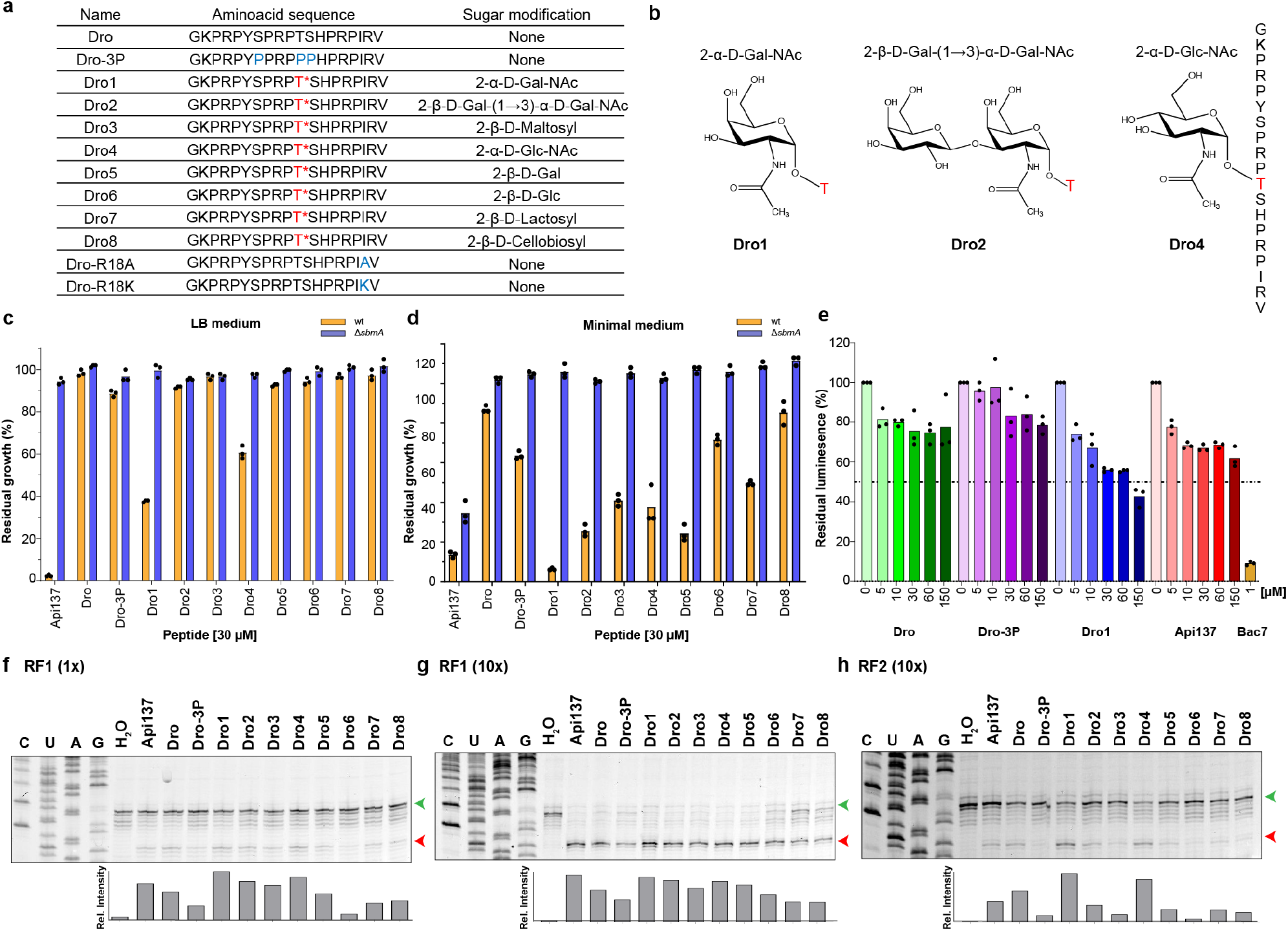
Characterization of inhibitory activity of drosocin derivatives. **a**, Amino acid sequences of the drosocin peptides used in this study. Drosocin peptides carrying a modification on Thr11 are indicated with T*, whereas the mutated positions are shown in blue. **b**, Chemical structures of the Thr11 modifications of Dro1, Dro2 and Dro4. **c**-**d**, *In vivo* inhibitory activity of 30 μM Api137 and drosocin derivatives on the growth of *E. coli* wt (yellow) and Δ*sbmA* (blue) strains in rich LB (**c**) or minimal medium (**d**). Histograms represent the averages from three biological replicates, individually plotted as dots. **e**, Inhibitory activity of increasing concentrations of Api137 (red), Dro (green), Dro-3P (purple), Dro1 (blue) and 1 μM Bac7 (gold) on *in vitro* translation using firefly luciferase as a reporter. The luminescence in the absence of compounds was normalized to 100 %; experiments were performed in triplicate and the bars represent the mean. **f**-**h**, Toeprinting assays monitoring the position of ribosomes on an MLIF*-mRNA in the presence of 30 μM Api137 and drosocin derivatives and either 1x RF1 (**f**), 10x RF1 (**g**) or 10x RF2 (**h**). Bands corresponding to ribosomes present at the start and stop codons are indicated by green and red arrows, respectively. The histogram represents the proportion of relative intensity of stop codon band for the different peptides.

While drosocin has been shown to inhibit protein synthesis *in vivo* and *in vitro* (Lele et al., 2015b; Ludwig et al., 2022), the exact mechanism by which it does so remains unclear. Interestingly, the type I insect PrAMP pyrrhocoricin is O-glycosylated with the same sugar at the same position of the peptide as drosocin, i.e. N-acetylgalactosamine on Thr11, and a minor disaccharide form with the additional galactose has also been detected (Cociancich et al., 1994). Together with the reported sequence similarity, drosocin was proposed to act analogously to the type I PrAMPs pyrrhocoricin and metalnikowins, rather than like the apidaecins and abaecins (Bulet et al., 1999). However, there are several subsequent observations that support similarity between drosocin and apidaecin, rather than type I PrAMPs. Firstly, in contrast to drosocin, unmodified pyrrhocoricin was shown to be slightly more active than the modified form (Hoffmann et al., 1999). Secondly, drosocin was suggested to belong to the apidaecin-like PrAMPs based on similarity in terms of ribosome-binding antibiotic competition assays, i.e. drosocin competes better with the type II PrAMP Api137 rather than the type I oncocin derivative Onc-112 (Krizsan et al., 2015). Lastly, drosocin lacking the carboxy-terminal Arg18-Val19 almost completely loses antimicrobial activity (Hoffmann et al., 1999), analogous to Api137 (Berthold and Hoffmann, 2014), whereas N-terminal rather than C-terminal truncations inactivate type I PrAMPs, such as Bac7 (Benincasa et al., 2004; Seefeldt et al., 2016).

Here we employ biochemical and structural approaches to dissect the mechanism by which drosocin interacts with the ribosome and inhibits protein synthesis, as well as shed light on the role of the critical O-glycosylation on Thr11. We show that the monosaccharide-modified drosocin is the most active, both in whole cell assays as well as within *in vitro* translation assays, suggesting that the modification is not only critical for cellular uptake, but also for ribosome binding and translation inhibition. In this regard, we demonstrate that the transporter SbmA plays a major role in uptake of drosocin, as reported for other PrAMPs (Florin et al., 2017; Mattiuzzo et al., 2007; Runti et al., 2013; Seefeldt et al., 2015). Moreover, we demonstrate that drosocin acts as a type II PrAMP, by interfering with translation termination, analogous to Api137, rather than acting during early elongation as a type I PrAMP. A cryo-electron microscopy (cryo-EM) structure of a drosocin-arrested ribosome at 2.3 Å, allowed the direct visualization of the complete drosocin peptide including the O-glycosylation. The structure reveals that drosocin has a completely different mode of interaction with the ribosome than Api137, and yet, like Api137, drosocin also utilizes a C-terminal arginine to directly interact and stabilize RF1 on the ribosome. Finally, we observe that the α-D-GalNAc modification on Thr11 of drosocin establishes multiple interactions with U2609 of the 23S rRNA, providing a structural basis for why glycosylation of drosocin peptides enhances the activity of drosocin peptides.

## Results

### SbmA plays a major role in drosocin uptake

Many PrAMPs, including Bac7, oncocin and apidaecin, utilize the SbmA transporter to pass through the *E. coli* inner membrane (Florin et al., 2017; Mattiuzzo et al., 2007; Runti et al., 2013; Seefeldt et al., 2015), however, whether drosocin also utilizes SbmA remains to our knowledge unknown. To address this, we monitored the effect of the presence of diverse drosocin peptides (**Fig. 1a,b**) on the growth of the wildtype *E. coli* strain BW25113 containing SbmA, as well as the *E. coli* BW25113 strain lacking SbmA (∆*sbmA*) (**Fig. 1c,d**). For our experiments, we compared unmodified drosocin (Dro) with various modified forms (Dro1-8) of drosocin (**Fig. 1a,b**). The modified forms included the naturally occurring Dro1 and Dro2 that carry either a monosaccharide (α-D-GalNAc) or disaccharide (β-D-Gal(1 →3)-α-D-GalNAc) attached to Thr11, respectively (**Fig. 1b**). In addition, we examined the previously reported (Lele et al., 2015a; Talat et al., 2011) drosocin derivatives bearing β-D-Maltosyl (Dro3), α-D-GlcNAc (Dro4), β-D-Gal (Dro5), β-D-Glc (Dro6), β-D-Lactosyl (Dro7) and β-D-Cellobiosyl (Dro8) modifications on Thr11 (**Fig. 1a,b**). Finally, we also included in our analysis the synthetic unmodified drosocin derivative with proline substitutions at positions 7, 11 and 12 (Dro-3P) (**Fig. 1a**), which was previously reported to have similar antimicrobial activity to the monosaccharide form of drosocin (Lele et al., 2015b). Growth was monitored in both rich (LB) and minimal medium in the presence of 30 µM of each peptide and normalized with the growth in the absence of the compounds (see Methods). In rich medium, we observe growth inhibition only with Dro1, bearing the monosaccharide α-D-GalNAc, and Dro4, which also carries a monosaccharide, namely, α-D-GlcNAc (**Fig. 1c**). Since no inhibition is observed with the monosaccharide β-D-Gal (Dro5) or β-D-Glc (Dro6) drosocins, this suggests that under these conditions the stereochemistry of the anomeric carbon on the sugar is more important than the type of sugar itself. We also observe no inhibition for Dro2 bearing the disaccharide β-D-Gal(1 →3)-α-D-GalNAc, nor for any of the β-linked disaccharides (Dro3, Dro6 or Dro7). Similarly, the unmodified drosocin and Dro-3P variant were also inactive in this assay (**Fig. 1c**). By contrast, all drosocin peptides inhibited growth of the *E. coli* BW25113 strain in minimal medium, albeit to different extents (**Fig. 1d**). The trends were similar to that reported previously (Lele et al., 2015a; Talat et al., 2011), namely, in that the highest inhibition was observed with Dro1 and the lowest with the unmodified peptide, whereas the other glycosylated variants lay in-between (**Fig. 1d**). We did not observe similar activity between Dro1 and Dro-3P as reported previously (Lele et al., 2015b), which may arise due to differences in the *E. coli* strains and/or growth conditions used. Strikingly, we note that any inhibition observed with the *E. coli* BW25113 strain was lost when performed with the BW25113 ∆*sbmA* strain, indicating that SbmA plays a major role in the cellular uptake of all drosocin peptides.

### Drosocin inhibits *in vitro* translation by trapping ribosomes at stop codons

Unmodified wildtype drosocin and Dro-3P peptides have been reported to inhibit *in vitro* translation reactions (Lele et al., 2015b; Ludwig et al., 2022), however, the naturally-occurring glycosylated form of drosocin has not been previously tested. To investigate this, we compared the effect of increasing concentrations (0-150 µM) of monosaccharide (α-D-GalNAc) modified (Dro1) with unmodified drosocin (Dro), Dro-3P and Api137 using a cell-free *in vitro* translation system with the firefly luciferase (Fluc) mRNA as a template (**Fig. 1e**), as we have used previously for assessing the activity of other PrAMPs (Mardirossian et al., 2018a; Mardirossian et al., 2018b; Mardirossian et al., 2019; Mardirossian et al., 2020; Seefeldt et al., 2016; Seefeldt et al., 2015; Sola et al., 2020). Dro1 exhibited dose-dependent inhibition, with an IC_50_ of 78 µM and reaching a maximum of 60% inhibition at the highest concentration tested of 150 µM. By contrast, both Dro and Dro-3P were poor inhibitors, reaching a maximum of 20% inhibition at 150 µM, whereas Api137 was slightly more effective, with 40% inhibition observed at 150 µM. This contrasts with type I PrAMPs, such as Bac7 (**Fig. 1e**) and Onc112, that display IC_50_ of <1 µM using the same system (Seefeldt et al., 2016; Seefeldt et al., 2015), suggesting that drosocin may inhibit translation similarly to Api137, rather than oncocin, as proposed previously (Krizsan et al., 2015).

To ascertain which step during protein synthesis is affected by drosocin, we performed toeprinting assays, where reverse transcription is used to monitor the position of ribosomes on a defined mRNA (Hartz et al., 1988). In the absence of PrAMP but presence of RF1, we observed no band corresponding to ribosomes at the UAA stop codon of the mRNA, whereas in the presence of 25 µM Api137 and RF1, ribosomes become stuck at the stop codon (**Fig. 1f** and **Supplementary Fig. 1**), as expected (Florin et al., 2017). Similarly, the same band was also observed in the presence of 30 µM of each of the tested drosocin derivatives, albeit with differing intensities (**Fig. 1f**). Increasing the concentration of RF1 by 10-fold in the reactions led to more intense termination bands (**Fig. 1g**), consistent with a role of drosocin acting during the termination phase, as reported for Api137 (Florin et al., 2017; Graf et al., 2018). We performed the same toeprinting reactions in the presence of 10-fold RF2, rather than 10-fold RF1, and also observed stalling of ribosomes at the stop codon, albeit with much lower efficiency (**Fig. 1h**). The strongest stalling was observed in presence of Dro1 and to a lesser extent with Dro4, a trend that was particularly evident in the presence of 10-fold RF2 (**Fig. 1h**). Both Dro1 and Dro4 were also the most active in our whole cells assays (**Fig. 1c-d**). By contrast, weak stalling was observed with Dro-3P, consistent with the lack of activity in the whole cell (**Fig. 1c,d**) and *in vitro* translation assays (**Fig. 1e**). Interestingly, we observed good activity for the unmodified drosocin peptide in the toeprinting assay (**Fig. 1f-h**), suggesting that the poor activity observed in the whole cell assays (**Fig. 1c,d**) may be due to cellular uptake. Collectively, our findings suggest that drosocin also traps ribosomes during termination, similar to the type II PrAMP apidaecin (Florin et al., 2017), but unlike the type I PrAMPs, such as Bac7 and Onc112 (Seefeldt et al., 2016; Seefeldt et al., 2015). Moreover, our results suggest that the monosaccharide-modified drosocin (Dro1) has the highest activity and that the glycosylation plays a role in cellular uptake, as well as binding of drosocin to the ribosome.

### Cryo-EM structures of drosocin-bound ribosome complexes

To investigate how drosocin inhibits translation and to provide insight into the role of the O-glycosylation, we set out to determine a cryo-EM structure of a ribosome-drosocin complex. Rather than forming complexes with vacant ribosomes or pre-defined functional states, we instead performed translation reactions with the same mRNA template used for the toeprinting assays in the presence of 10-fold RF1 and 30 µM Dro1 (**Fig. 1g**). Reactions were subsequently pelleted through sucrose cushions and the pelleted ribosomal complexes were subjected to single particle cryo-EM analysis. *In silico* sorting of the data revealed three main populations of ribosomal states, namely, 70S ribosomes with RF1 and P-site tRNA (26.0%), or with A- and P-site tRNAs (16.0%), as well as a population containing only large 50S subunits (30.2%) (**Supplementary Fig. 2**), which after refinement yielded final reconstructions at 2.3 Å, 2.8 Å and 2.1 Å, respectively (**Fig. 2a-c** and **Supplementary Fig. 3**). In all three reconstructions, additional density was observed within the ribosomal exit tunnel that could be unambiguously assigned to the drosocin peptide (**Fig. 2a-i**). The density for drosocin was particularly well-resolved in the RF1-containing 70S map enabling all 19 amino acids to be modelled with sidechains (**Fig. 2d** and **Supplementary Fig. 3**), including the α-D-GalNAc modification linked to Thr11 (**Fig. 2g**). Similarly, the density for drosocin in the cryo-EM map of the 50S subunit was generally well-resolved, except for the N- and C-terminal regions (**Fig. 2f,i** and **Supplementary Fig. 3**). By contrast, the density for drosocin in the cryo-EM map of the complex containing A- and P-site tRNAs was less well-resolved (**Fig. 2e** and **Supplementary Fig. 3**), and was particularly poor for the α-D-GalNAc modification (**Fig. 2h**). This suggested to us that the peptide is bound less stably within this complex. Nevertheless, in all three structures, the overall orientation of the drosocin peptide within the exit tunnel was identical, namely, with the C-terminus located at PTC and the N-terminus extending into exit tunnel, analogous to an elongating nascent polypeptide chain (**Fig. 2a-f** and **Supplementary Fig. 4a**). This orientation is also the same as that observed for the type II PrAMP Api137 (Chan et al., 2020; Florin et al., 2017; Graf et al., 2018) and opposite to that of type I PrAMPs, such as Bac7 and pyrrochoricin (**Supplementary Fig. 4b-d**) (Gagnon et al., 2016; Mardirossian et al., 2018b; Mardirossian et al., 2020; Roy et al., 2015; Seefeldt et al., 2016; Seefeldt et al., 2015).

**Fig. 2:**
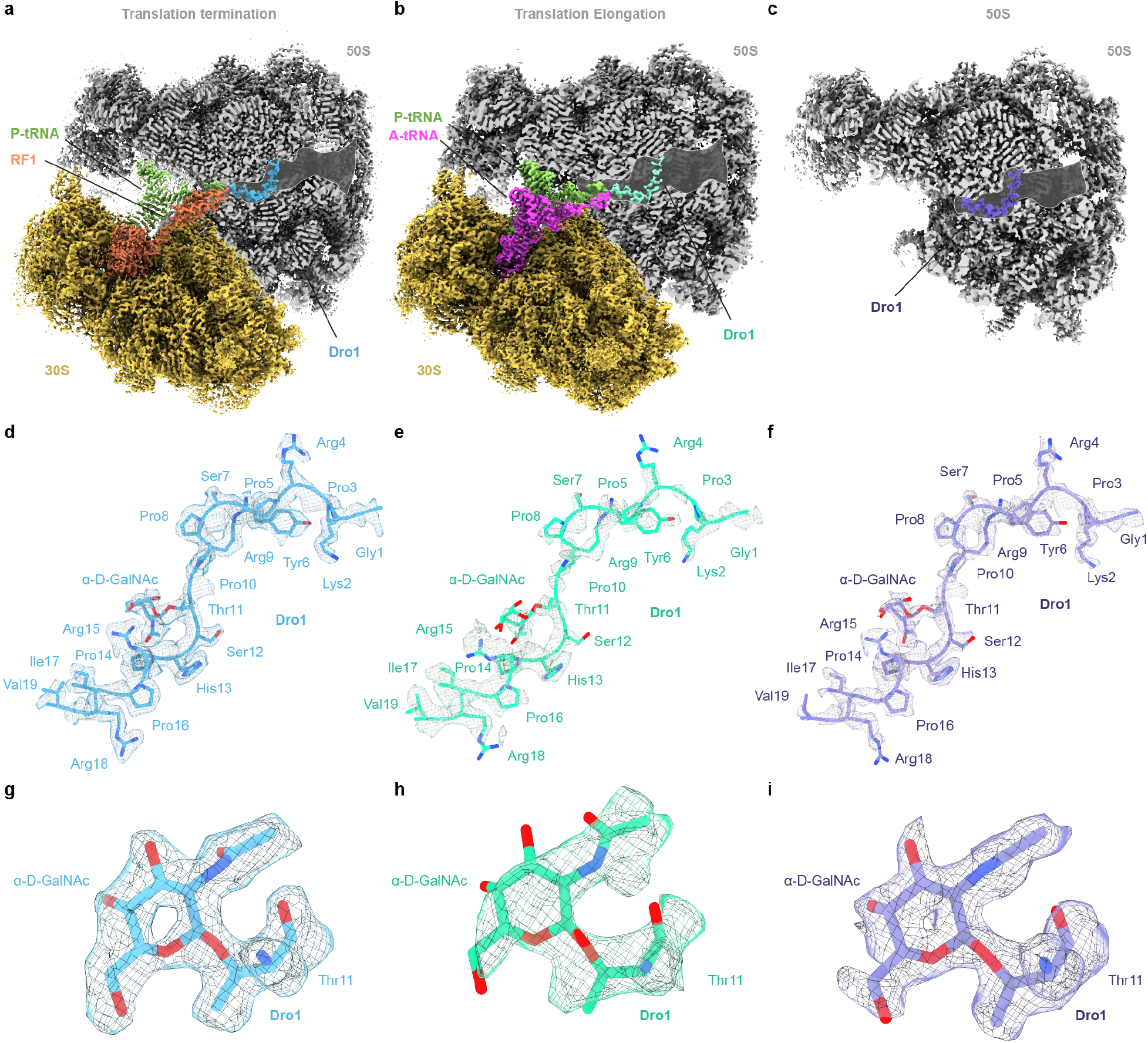
Cryo-EM structures of drosocin-bound ribosomal complexes. **a-c**, Cryo-EM maps of Dro1-bound to (**a**) termination and (**b**) elongation complexes, as well as the (**c**) large 50S subunit, with transverse section of the 50S (grey) to reveal the Dro1 binding site within the exit tunnel. In (**a**) the P-tRNA, RF1 and Dro1 are coloured green, orange and cyan, respectively. In (**b**) the A-tRNA, P-tRNA and Dro1 are coloured green, pink and teal, respectively, whereas in (**c**) Dro1 is purple. **d-f**, Cryo-EM density (grey mesh) with molecular model for (**d**) Dro1 (cyan) from termination complex as in (**a**), (**e**) Dro1 (teal) from elongation complex as in (**b**), and (**f**) Dro1 (purple) from the 50S subunit as in (**c**). **g-i**, Cryo-EM density (grey mesh) with molecular model for α-D-GalNAc modification at Thr11 of Dro1 in (**g**) the termination and (**h**) elongation complexes, as well as (**i**) the 50S subunit.

### Cryo-EM structure of drosocin bound to an elongating ribosome

For the drosocin-ribosome complex containing A- and P-site tRNAs, comparison of the cryo-EM density (**Fig. 3a**) with pre- and post-attack states (Polikanov et al., 2014) (**Fig. 3b-c**) indicates that the P-site tRNA is deacylated, whereas the A-site tRNA carries a nascent chain (**Fig. 3a**). Thus, drosocin is bound to an elongating ribosome state that is post-peptide bond formation, but pre-translocation. Inspection of the cryo-EM density for the anticodon-codon interactions suggests that the A- and P-site tRNAs contain initiator tRNA^fMet^ and tRNA^Leu^ decoding the AUG and UUC codons in the first and second positions of the mRNA, respectively (**Supplementary Fig. 5a,b**), and therefore the nascent chain should comprise the dipeptide fMet-Leu. This would also be consistent with the limited space available at the PTC for the nascent chain due to the presence of drosocin blocking the ribosomal exit tunnel. However, because the density for the nascent chain is poorly resolved and thus could not be modelled *de novo*, we could only generate a tentative model for fMet-Leu, nevertheless illustrating that the position is different than for fMet-Phe in the post-peptide bond formation state reported previously (**Fig. 3c,d**)(Polikanov et al., 2014). In the latter, we would predict steric clashes between the fMet moiety and the N-terminal Val19 of drosocin (**Fig. 3c**), which appears to have forced the fMet moiety to shift towards Arg18 (**Fig. 3d**), providing a likely explanation as to why both regions are poorly ordered in this complex (**Fig. 3a**). Collectively, these findings suggest that for this elongating complex to exist, drosocin permits initiation (despite predicted clashes between the fMet and Val19 as seen in **Fig. 3b**), aminoacyl-tRNA binding to the A-site and subsequent peptide bond formation, but interferes with the first translocation step. In order to mimic the translocated state, we modelled fMet-Leu-tRNA bound in the P-site based on other available P-site peptidyl-tRNAs (Syroegin et al., 2022a), which revealed even larger steric clashes with drosocin (**Fig. 3e**), providing a structural explanation for the observed translocation inhibition. We note that while apidaecin strongly interferes with termination, moderate effects on initiation have also been reported *in vivo* and *in vitro* (Mangano et al., 2020). Given the similarity in the binding position of the C-terminus of Api137 and drosocin on the ribosome (**Supplementary Fig. 4b**), it seems likely that apidaecin may also interfere with the first translocation step as seen here for drosocin, rather than acting like a type I PrAMP to prevent accommodation of the aminoacyl-tRNA at the A-site of the PTC, but this remains to be determined.

**Fig. 3:**
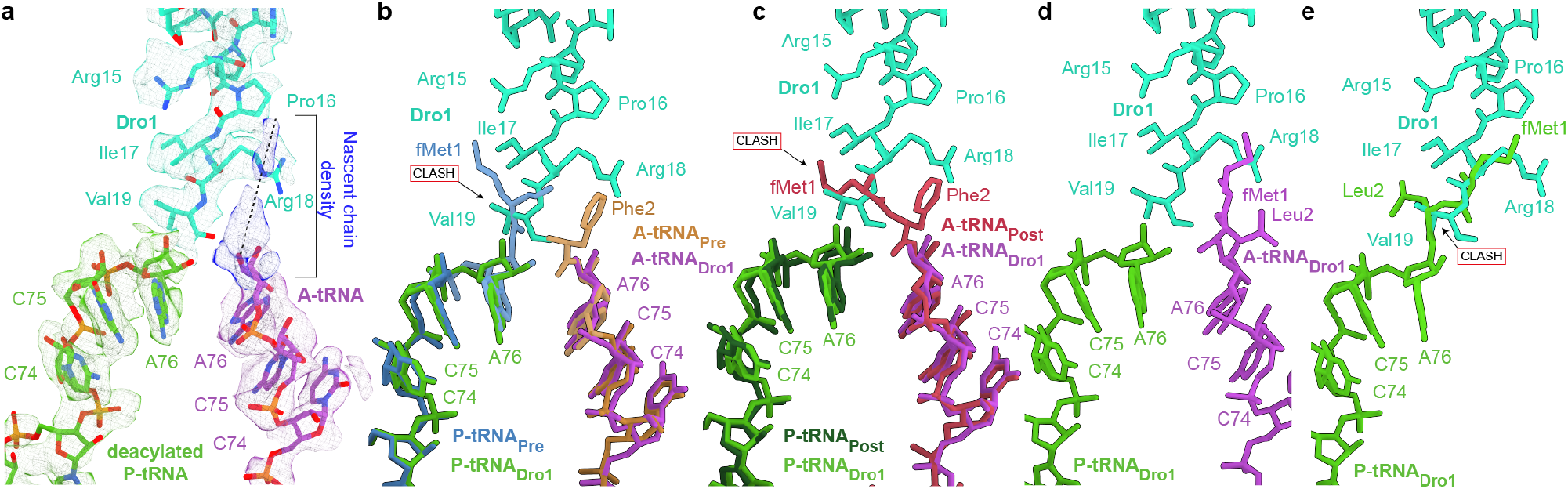
Cryo-EM structure of the drosocin-bound translation elongation complex. **a**, Isolated cryo-EM densities (mesh) with molecular models for P-tRNA (light green), A-tRNA (magenta) and Dro1 (teal) within the translation elongation complex. Additional density connected to the A-site tRNA is attributed to the nascent chain, but cannot be modelled due to flexibility. **b**, Superimposition of P-tRNA_Dro1_ (light green), A-tRNA_Dro1_ (purple) and Dro1 (teal) from (**a**), with P-tRNA_Pre_ (blue) and A-tRNA_Post_ (brown) from PRE-state (PDB ID 1VY4) (Polikanov et al., 2014). Alignment based on the 23S rRNA. The fMet attached to the P-site tRNA would be predicted to clash with the C-terminus of Dro1 (teal). **c**, Superimposition of P-tRNA_Dro1_ (light green), A-tRNA_Dro1_ (purple) and Dro1 (teal) from (**a**), with P-tRNA_Post_ (dark green) and A-tRNA_Post_ (red) from POST-state (PDB ID 1VY5) (Polikanov et al., 2014). Alignment based on the 23S rRNA. The fMet from the fMet-Phe, attached to the A-tRNA Post would be predicted to clash with the C-terminus of Dro1 (teal). **d**, Hypothetical molecular model of the fMet-Leu nascent chain connected to the A-tRNA (based on POST-state PDB ID 1VY5) (Polikanov et al., 2014). **e**, Steric clash of the fMet-Leu nascent chain in the P-site after translocation (based on PDB ID 7RQE) (Syroegin et al., 2022b).

### Interaction of drosocin within the tunnel of the RF1-bound complex

In the RF1-bound complex, Dro1 is very well-resolved enabling a molecular description of the interactions of the drosocin peptide with components of the ribosomal tunnel as well as RF1 (**Fig. 4**). The N-terminus of Dro1 reaches down the tunnel past the constriction created by the extensions of ribosomal proteins uL4 and uL22 (**Fig. 4a**), where the N-terminal amino group can form potential hydrogen bonding interactions with the N7 of 23S rRNA nucleotide G1259 (**Fig. 4b**). Additionally, Lys2 establishes two contacts with U1258, one from the backbone amine to the phosphate-oxygen of U1258, and the other mediated via a water molecule between the ε-amino group of the Lys2 sidechain and the O4 of U1258 (**Fig. 4b**). Removal of first five N-terminal residues (GKPRP), which includes the first PRP motif, completely abolishes activity, suggesting the importance of the N-terminal interactions for drosocin activity, although effects on uptake cannot be excluded (Bulet et al., 1996). Residues Pro3 to Pro10 of Dro1 are located at the constriction and establish multiple interactions with uL4 and uL22 (**Fig. 4c-e**). Specifically, the backbone carboxyls of Pro3 and Tyr6 of Dro1 are within hydrogen bonding distance to the sidechains Arg67 and Arg61 of uL4, respectively (**Fig. 4c**). Interactions with uL22 include hydrogen bonds between the sidechains of Tyr6 and Arg9 of Dro1 with the backbone of Ala93 and Lys90/Gly91 of uL22, respectively (**Fig. 4d**). The Tyr6 interaction appears not to be critical since mutation to Phe that lacks the hydroxyl group does not lead to loss of antimicrobial activity (de Visser et al., 2005). In addition, the backbone carboxyl of Pro10 of Dro1 can interact with the sidechain of Lys90 of uL22 as well as indirectly with Arg92 via a water molecule (**Fig. 4e**). Mutation of Pro10 to Ala abolishes antimicrobial activity (Ahn et al., 2011b), presumably by altering the conformation of the peptide within this region. Ser12 of Dro1 can hydrogen bond directly with U746 and form a water-mediated interaction with G748 (**Fig. 4e**). The α-D-GalNAc modification on Thr11 establishes multiple interactions with U2609, which is discussed in more detail in a following section.

**Fig. 4:**
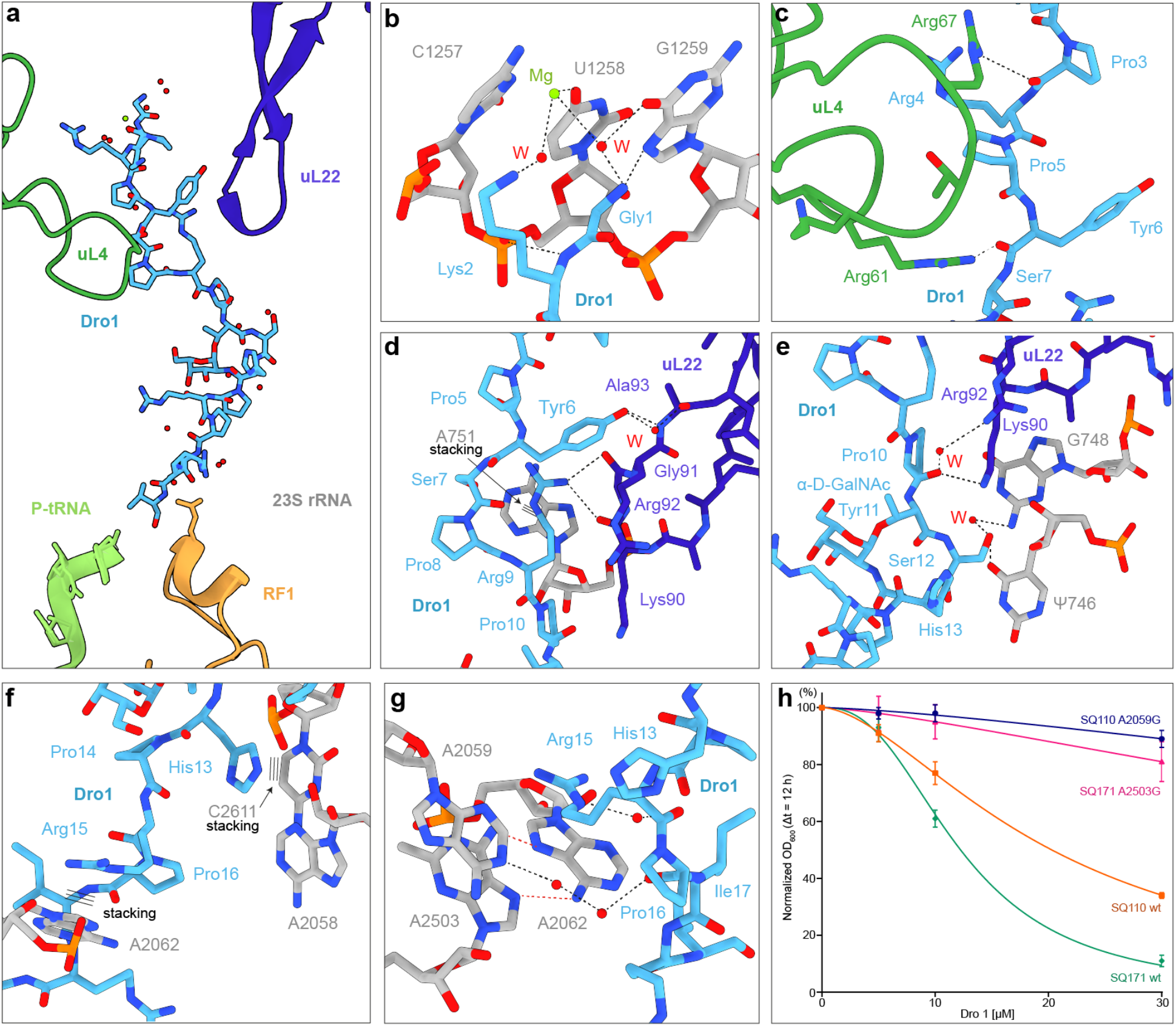
Interactions of drosocin within the exit tunnel. **a-g**, Dro1 (light blue) in the nascent peptide exit tunnel (NPET) with surrounding 23S rRNA nucleotides (grey), with P-tRNA (lime), RF1 (orange), uL4 (green) and uL22 (dark blue). **b**, Water-mediated and direct hydrogen bond interactions of Gly1 and Lys2 of Dro1 with U1258 and G1259. **c**, Hydrogen bond interactions of Arg61 and Arg67 of uL4 with the backbone of Pro3 and Tyr6 of Dro1. **d**, Stacking interaction (indicated by three lines) of A751 with Arg9 and water-mediated and direct interactions of Tyr6 and Arg9 of Dro1 with Lys90, Gly91, Arg92 and Arg93 of uL22. **e**, Water-mediated and direct interactions of Pro10 backbone and Ser12 of Dro1 with Lys90 and Arg92 of uL22 and with G748 and Ψ746. **f**, Stacking interactions of His13 and Arg15 with C2611 and A2062 respectively (stacking indicated by three lines). **g**, Water-mediated and direct hydrogen bond interactions of Arg15 and Pro16 of Dro1 with the backbone with A2059, A2062 and A2503. **h**, *in vivo* inhibitory activity of 5 μM, 10 μM and 30 μM Dro1 on the growth of *E. coli* SQ110 wt (orange), *E. coli* SQ110 A2059G (blue), *E. coli* SQ171 wt (green) and *E. coli* SQ171 A2503G (pink) in LB medium. For each concentration, residual growth values are the OD_600_ at t = 12 h of the treated culture normalized to the untreated one, considered as 100 %. Error bars represent the standard deviation for three biological replicas and the measurement error of the plate reader. The curves were calculated and plotted by non-linear regression.

### Stacking interactions and drosocin resistance mutations

In total, there are three stacking interactions observed between sidechains of Dro1 and nucleobases of the 23S rRNA, namely, between Arg9 and A751 (**Fig. 4d**), His13 and C2611, as well as Arg15 and A2062 (**Fig. 4f,g**). Mutation of Arg9 or Arg15 to lysine reduces antimicrobial activity of the Dro peptides by 4- and 8-fold, respectively (Lele et al., 2013), suggesting that these interactions contribute to drosocin binding. Api137 also establishes stacking interactions with A751 and C2611 (Chan et al., 2020; Florin et al., 2017; Graf et al., 2018), however, the sidechains and modes of interaction are completely distinct (**Supplementary Fig. 6a-d**). Compared to the canonical RF1-bound termination complexes (Fu et al., 2019; Laurberg et al., 2008; Pierson et al., 2016; Zhou et al., 2012), we observe a rotated conformation of A2062 (**Fig. 4g** and **Supplementary Fig. 6e-g**), which is also observed in the Api137-bound ribosome structures (Chan et al., 2020; Florin et al., 2017; Graf et al., 2018) (**Supplementary Fig. 6e-g**). The rotated conformation of A2062 forms interactions with A2503, which is adjacent to A2059 (**Fig. 4g**), both of which were shown to confer resistance to Api137 when mutated (Florin et al., 2017). Since Arg15 of Dro1 stacks upon A2062 (**Fig. 4g**), and is in close proximity of A2503 and A2059, we assessed whether A2503G and A2059G mutations confer resistance to Dro1. Indeed, we observed that compared to the wildtype strain, both strains bearing the A2503G and A2059G mutations were more resistant to Api137 (**Supplementary Fig. 6h**), as previously reported (Florin et al., 2017), but also to Dro1 (**Fig. 4h**). We believe these findings provide strong evidence that the ribosome (and therefore translation) is a (if not “the”) physiological target for Dro1 within the bacterial cell. This is also supported by the identification of mutations in ribosomal protein uL16 and RF2 that confer resistance to Api137, also confer resistance to Dro (see accompanying manuscript of *Mangano* et al 2022).

### C-terminal interactions are critical for drosocin activity

The C-terminus of Dro1 is stabilized by three backbone interactions between residues Ile17-Arg18 and the bases of 23S rRNA nucleotides U2506, G2061 and A2062 (**Fig. 5a**).

**Fig. 5:**
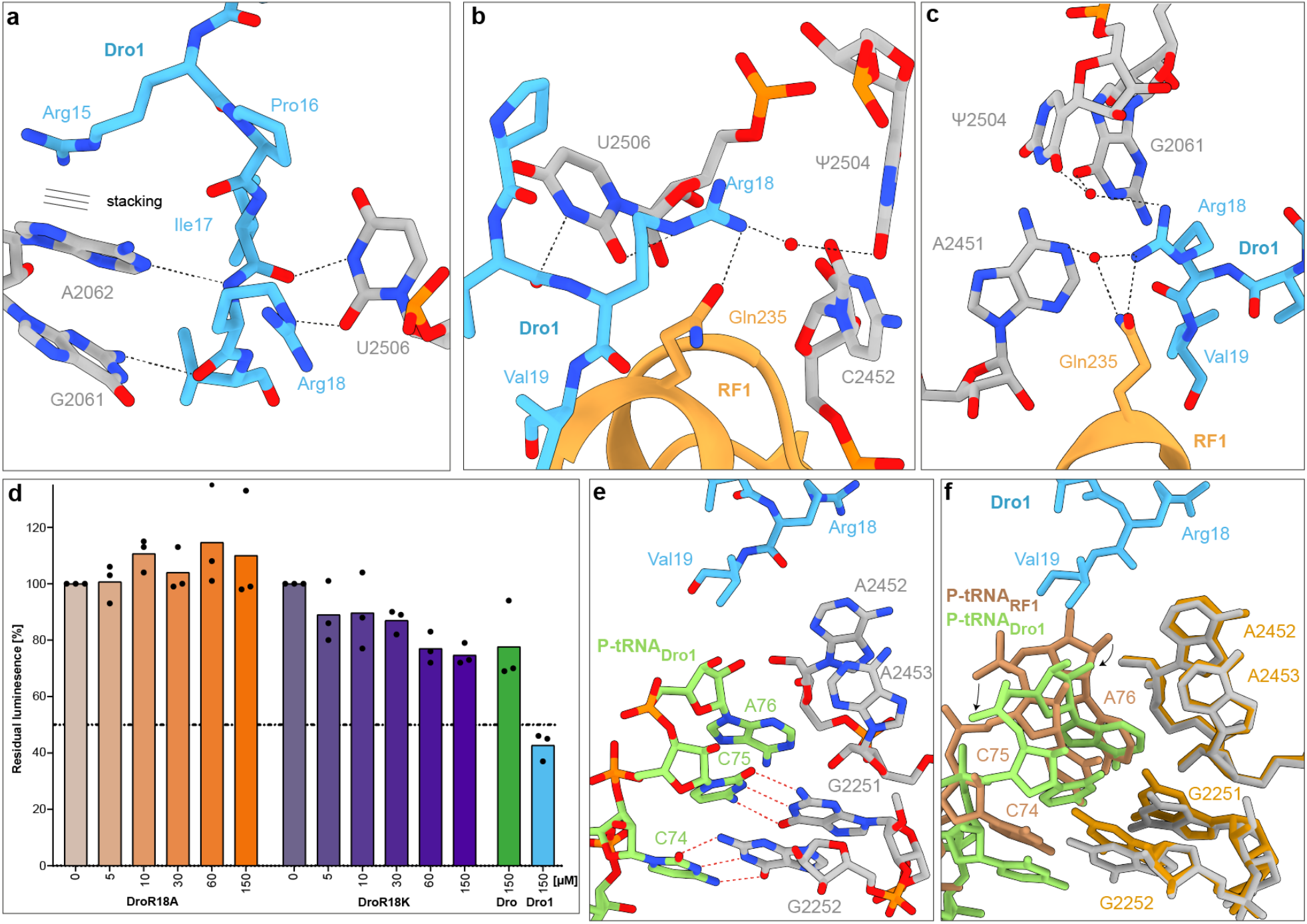
Interactions of drosocin with RF1 and P-tRNA. **a-c**, Interactions of Dro1 (light blue) with 23S rRNA nucleotides (grey) and RF1 (orange). **a**, Stacking interaction of Arg15 (indicated by three lines) and hydrogen bond interactions of G2061, A2062 and U2506 with Ile17 and Arg18 of Dro1. **b-c**, Two views of the water-mediated and direct hydrogen bond interactions of Arg18 of Dro1 with Gln235 of RF1 and 23S rRNA nucleotides (**b**) Ψ2504, U2506 and C2452 and (**c**) G2061, A2451 and Ψ2504. **d**, Inhibitory activity of increasing concentrations of Dro R18A (orange), Dro R18K (purple), and 150 μM Dro (green) or Dro1 (blue), on *in vitro* translation using firefly luciferase as a reporter. The luminescence in the absence of compounds was normalized to 100 %; experiments were performed in triplicate and the bars represent the mean. **e**, Deacylated P-tRNA (lime) in the presence of Dro1 (light blue). **f**, superimposition of (**e**) with a P-tRNA from a canonical termination complex (brown, PDB ID 4V63) (Laurberg et al., 2008). Dro1 displaces the CCA-end of the deacylated tRNA while keeping the base pairing interactions of C74 and C75 with G2252 and G2251 (grey) respectively which slightly tilts the nucleotides compared to the canonical position (yellow).

Additionally, the sidechain of Arg18 inserts into a pocket where it can form direct hydrogen bonds with the nucleobases of C2452 and U2506 (**Fig. 5b**), as well as via water-mediated interactions with U2504, G2061 and A2451 (**Fig. 5b,c**). Importantly, Arg18 comes within 2.9 Å of Gln235 of the conserved GGQ motif of RF1, and a further water-mediated interaction with Gln235 is also possible (**Fig. 5b,c**), suggesting Arg18 plays an important role in stabilizing RF1 on the ribosome. This interaction is reminiscent of that observed previously between Arg17 of Api137 and Gln235 of RF1 (Chan et al., 2020; Florin et al., 2017; Graf et al., 2018) (**Supplementary Fig. 7a-f**), the importance of which was shown by Arg17Ala mutations that decrease both the ribosome affinity and inhibitory activity of the peptide (Krizsan et al., 2014). While deletion of the last two residues (Arg18-Val19) of drosocin completely abolished *in vitro* biological activity (Hoffmann et al., 1999), single substitutions of Arg18 have to our knowledge not been undertaken. Therefore, we synthesized an unmodified drosocin peptide bearing the Arg18Ala mutation (**Fig. 1a**) and tested its activity using *in vitro* translation assays, demonstrating a complete loss of activity for the Dro-R18A peptide (**Fig. 5d**). By contrast, Dro bearing an Arg18Lys mutation (Dro-R18K, **Fig. 1a**) displayed similar activity to the unmodified wildtype Dro peptide (**Fig. 5d**). Unlike Arg18, the very C-terminal Val19 of Dro1 is poorly ordered in the complex, but at lower thresholds density is observed to encroach on the binding site of a canonical P-site tRNA located at the PTC (**Fig. 5e,f**). As a consequence, the CCA-end of the P-site tRNA, which is also poorly resolved, is clearly shifted by 2-3 Å from its canonical position observed in RF1-termination complexes (Fu et al., 2019; Laurberg et al., 2008; Pierson et al., 2016; Zhou et al., 2012) (**Fig. 5e,f**). The shift is predominantly of the backbone of the CCA-end enabling the nucleobases of C74 and C75 to maintain Watson-Crick base-pairs with P-loop nucleotides G2252 and G2251, respectively (**Fig. 5e,f**). This is distinct from Api137, where the C-terminus was observed to directly interact with the A76 of the P-site tRNA and stabilize the P-site tRNA in its canonical position (**Supplementary Fig. 7g-h**). By comparison, we do not observe a shifted P-site tRNA in the Dro1-bound elongating state (**Supplementary Fig. 7i**). Otherwise, the binding position and interactions of RF1 in the Dro1-RF1-ribosome complex are identical to those observed previously for RF1 decoding of stop codons during canonical termination (Fu et al., 2019; Laurberg et al., 2008; Pierson et al., 2016; Zhou et al., 2012) (**Supplementary Fig. 8a-d**). However, with the higher resolution we also observe multiple water-mediated interactions between RF1 and the UAA stop codon (**Supplementary Fig. 8a-d**), which were not reported in the previous lower resolution termination complexes (Fu et al., 2019; Laurberg et al., 2008; Pierson et al., 2016; Zhou et al., 2012).

### Interaction of the O-glycosylation of drosocin with the ribosome

For the Dro1-RF1-70S and Dro1-50S complexes, the α-D-GalNAc modification linked to Thr11 establishes multiple interactions with U2609 of the 23S rRNA (**Fig. 6a**). In particular, the C3 hydroxyl comes within 2.6 Å and 2.7 Å of the N3 and O2, respectively, of the base of U2609 (**Fig. 6a**). Additionally, a hydrogen bond is also possible (3.5 Å) from the C4 hydroxyl to the O4 of U2609 (**Fig. 6a**). We note that the α-D-GlcNAc modification present in Dro4 would maintain the former interactions, and only lose the latter weaker interaction with O4 of U2609 (**Supplementary Fig. 9a,b**), consistent with the similar activity of Dro4 compared to Dro1 (**Fig. 1c,d**). By contrast, modifications of β-D-linkage as in Dro3 and Dro5-Dro8, would be incompatible with the interactions observed for the α-D-GlcNAc modification, providing an explanation why they exhibit lower activity compared to Dro1 and Dro4 (**Fig. 1c,d**). Comparison with other *E. coli* 70S ribosome structures, including RF1-termination complexes (Fu et al., 2019; Laurberg et al., 2008; Pierson et al., 2016; Zhou et al., 2012), reveals that U2609 is usually base-paired with A752 (**Fig. 6b and Supplementary Fig. 9c**), whereas in the Dro1-RF1-70S and Dro1-50S complexes, the α-D-GalNAc modification occupies the position of U2609, causing the base to shift away from A752 by up to 6 Å (**Fig. 6b,c** and **Supplementary Fig. 9d,e**). Moreover, we observe two waters molecules located between U2609 and A752 that may also contribute to stabilizing the shifted conformation by establishing indirect interactions between U2609 and the α-D-GalNAc modification of Dro1 (**Fig. 6a,c**). Interestingly, in the cryo-EM map of Dro1 bound to the elongating ribosome, we observe both the base-paired and shifted conformation of U2609 (**Fig. 6d** and **Supplementary Fig. 9f**). As mentioned, the density for the α-D-GalNAc modification of Dro1 is less well-resolved in this complex (**Fig. 2h** and **Supplementary Fig. 9f**), suggesting that it is highly flexible, presumably because it cannot adopt the preferred position interacting with the shifted conformation of U2609.

**Fig. 6:**
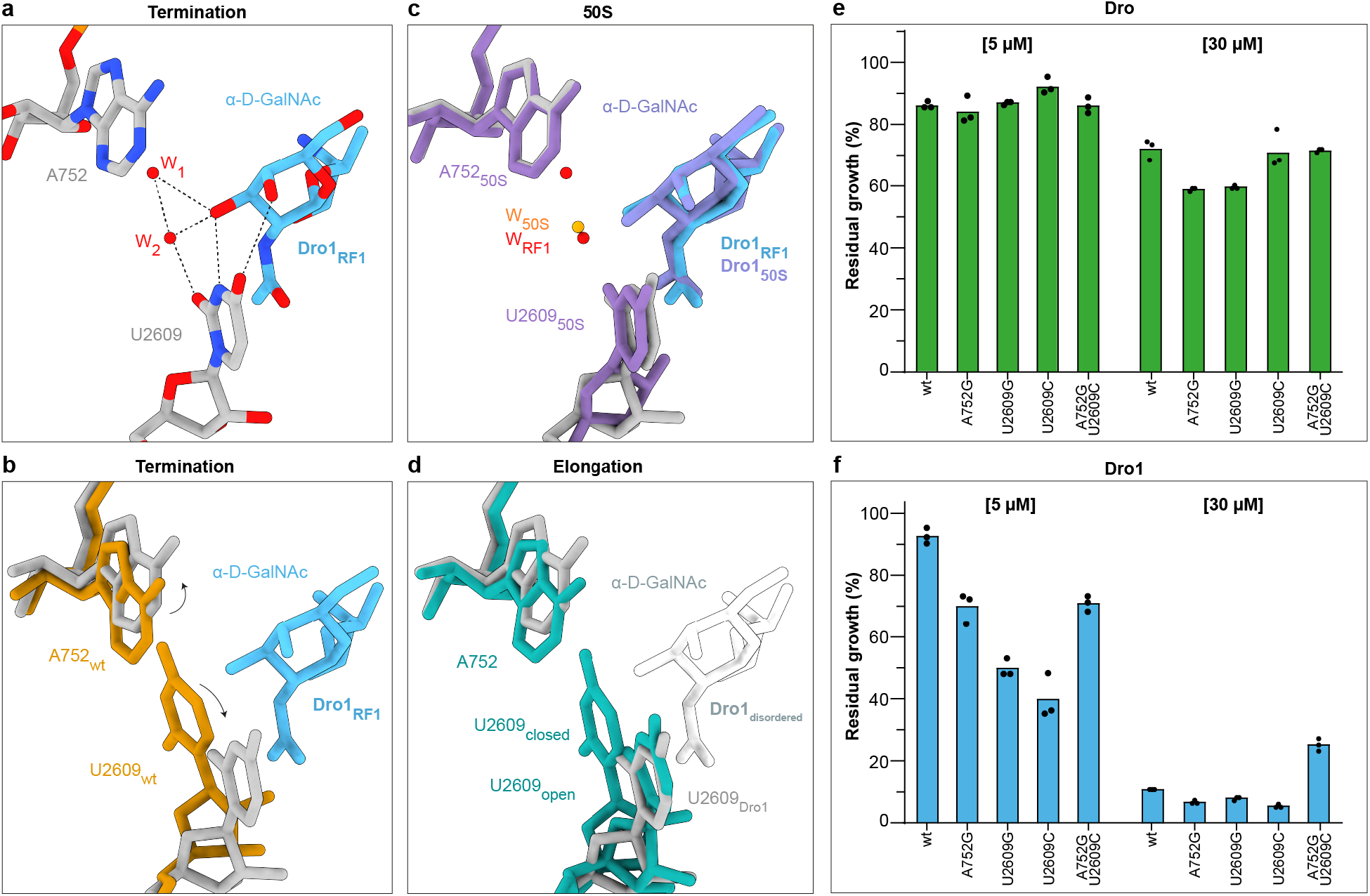
Interaction of O-glycosylation of Dro1 with U2609 of the 23S rRNA. **a**, Molecular interactions between the α-D-GalNAc modification on Thr11 of Dro1 (light blue) and the 23S rRNA nucleotide U2609 (grey) of the Dro1-bound termination complex. Two coordinated water molecules (red) stabilize the interactions of α-D-GalNAc of Dro1 with U2609. **b**-**d**, Superimposition of Dro1_RF1_ (light blue), waters (red) and 23S rRNA (grey) from (**a**) with (**b**) 23S rRNA (yellow) from canonical RF1-bound termination complex (PDB ID 4V63) (Laurberg et al., 2008), (**c**) 23S rRNA (purple) from the Dro1-bound 50S complex and (**d**) 23S rRNA (turquoise) from the Dro1-bound elongation complex with two alternative conformations (open and closed) of U2609 shown. α-D-GalNAc modification of Dro1 was poorly ordered in the elongation complex, therefore, the white silhouette indicates the position from Dro1_RF1_ that is incompatible with the closed conformation of U2609. **e-f**, *In vivo* inhibitory activity of 5 μM and 30 μM of (**e**) Dro (green) and (**f**) Dro1 (blue) on the growth of *E. coli* SQ171 wt, *E. coli* SQ171 A752G, *E. coli* SQ171 U2609G, *E. coli* SQ171 U2609C and *E. coli* SQ171 A752G/U2609C. Histograms represent the averages from three biological replicates, individually plotted as dots.

**Fig. 7:**
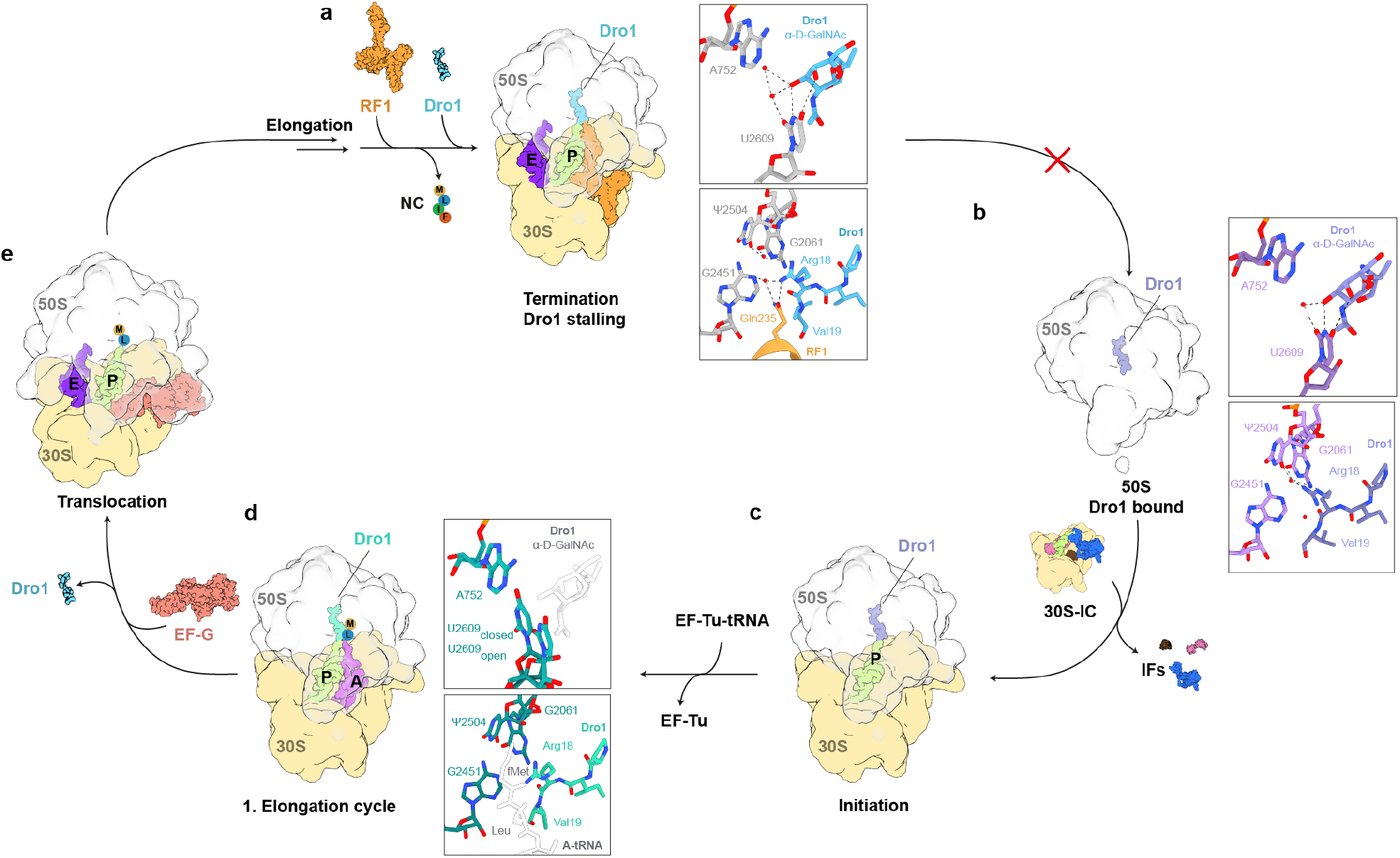
Model for the mechanism of action of Dro1 inhibition during translation. **a**, Appearance of a stop codon in the A-site is recognized by RF1 (or RF2, orange), which catalyzes release of the nascent chain (NC) from the P-site tRNA (lime). Following NC release, Dro1 (light blue) binds within the exit tunnel, separating the A752-U2609-basepair (grey) with Dro1 α-D-GalNAc modification and becomes stabilized via water-mediated and direct interactions between Arg18 of Dro1 and the Gln235 of the conserved GGQ motif of RF1 and surrounding 23S rRNA nucelotides. This interaction stabilizes RF1 on the post-release complex, preventing its dissociation and thereby blocking subsequent ribosome recycling steps and re-initiation. **b**, Dro1 (purple) binds to free 50S subunits (grey), separating the A752-U2609-basepair (light purple) with Dro1 α-D-GalNAc modification but is not fully stabilized via water-mediated and direct interactions between Arg18 of Dro1 and surrounding 23S rRNA nucleotides. **c**, Translation initiation complexes can form in the presence of Dro1 (purple), despite slight overlap between Dro1 and the fMet moiety of the P-tRNA (lime), suggesting that fMet might displace the C-terminal part of Dro1. **d**, Following peptide bond formation, the presence of Dro1 (teal) appears to interfere with translocation of the dipeptidyl-tRNA in the A-site (purple) into the P-site (lime). The α-D-GalNAc modification (white) is disordered and both the open and closed conformation of the U2609 base (dark teal) is observed. The dipeptidyl moiety (white) on the A-tRNA interferes with the stabilization of Dro1 in the PTC. **e**, For translocation to occur, and subsequent steps of elongation to occur, Dro1 must dissociate from the ribosome followed by elongation until translation termination is reached.

Collectively, these findings suggest that the propensity of the U2609-A752 to base-pair could influence the ability of Dro1 to bind stably to the ribosome and inhibit translation. To test this, we monitored the antimicrobial activity of Dro1 on strains bearing either A752G, U2609G or U2609C mutations, which should perturb Watson-Crick base-pairing. In addition, we also used a strain with a U2609C-A752G double mutation, which would be predicted to restore Watson-Crick base-pairing, and with three hydrogen bonds could possibly make breaking the base-pair harder than with the canonical two hydrogen bonds for the A-U base-pair. As a control, we also tested Dro that lacks the α-D-GalNAc modification, which we predict (assuming that Dro binds analogously to Dro1) should not interact with U2609 and therefore not be influenced by the conformation of the U2609-A752 base-pair. As seen in **Fig. 6e**, we observed that there was no significant difference in growth inhibition by 5 µM Dro, and only a modest effect at 30 µM Dro, when comparing the wildtype strain and strains bearing single or double mutations. By contrast, we observed that the growth of the strains bearing the single point mutations was more susceptible to Dro1 than the wildtype strain, especially for the U2609C mutation, although this effect became less evident at higher (30 µM) drug concentrations (**Fig. 6f**). Although the U2609C-A752G double mutation was also slightly more susceptible than the wildtype to Dro1 at 5 µM, it was still less susceptible than most single point mutations, and appeared to be 2.5-fold more tolerant to Dro1 than the wildtype strain at 30 µM (**Fig. 6f**). Collectively, these findings support a role for the U2609-A752 base-pair in modulating the ribosome binding and inhibition activity of glycosylated drosocin. Overall, there is excellent agreement between the interactions observed here for Dro1 and the extensive mutagenesis performed on Dro in the accompanying study of Mangano *et al* (2022).

## Discussion

Our biochemical and structural analysis allows us to propose a model for the mechanism of action of drosocin, highlighting the role of the O-glycosylation (**Fig. 7**). Analogous to Api137 (Chan et al., 2020; Florin et al., 2017; Graf et al., 2018), we reveal that drosocin interferes with the translation termination by trapping RF1 on the ribosome subsequent to the release of the nascent polypeptide chain (**Fig. 7a**). Like Api137 (Chan et al., 2020; Florin et al., 2017; Graf et al., 2018), an arginine residue (Arg18) at the C-terminus of the Dro directly interacts with Gln235 of the conserved GGQ motif of RF1 (**Fig. 7a**). Arg18 of Dro is critical since mutation to alanine abolishes all inhibitory activity of the peptide (**Fig. 5d**), collectively providing a structural basis for how RF1 dissociation is impeded by drosocin. Unlike Api137, drosocin is O-glycosylated on Thr11 and we observe that the α-D-GalNAc modification contributes to the ribosome binding by establishing multiple hydrogen bond interactions with U2609 of the 23S rRNA (**Fig. 7a**). This interaction rationalizes our (**Fig. 1**), and previous (Ahn et al., 2011a; Ahn et al., 2011b; Gobbo et al., 2002; Lele et al., 2015a; Marcaurelle et al., 1998; Otvos et al., 2000; Rodriguez et al., 1997; Talat et al., 2011) observations that the native modified forms of drosocin generally display enhanced antimicrobial activity compared to the unmodified peptide. Interestingly, we observe drosocin causes a shift of U2609 that breaks the base-pair that U2609 usually forms with A752 (**Fig. 7a**). Consistently, we could demonstrate that single and double mutations at these positions could influence the activity of the glycosylated, but not unmodified, form of drosocin (**Fig. 6e,f**). To our knowledge, breaking of this base-pair has not been observed in *E. coli* previously, although the base-pair has been shown to be important for interaction for ketolide antibiotics, such as telithromycin (Dunkle et al., 2010) and, in particular, for their bactericidal activity (Svetlov et al., 2020). We note however that U2609 and A752 are unpaired in some bacterial ribosomes, such as *Mycobacterium tuberculosis* (Yang et al., 2017), raising the question of whether these ribosomes are more susceptible to glycosylated forms of drosocin.

In addition to the termination complex, we observed drosocin bound to two other ribosomal particles, namely, a vacant 50S subunit and an elongating ribosome (**Fig. 2b,c**). This implies that in the cell, drosocin could potentially interact with the 50S subunit following termination and ribosome recycling, when the 70S ribosomes are split into their component subunits (**Fig. 7b**). This is not surprising given that the majority of the interactions formed by drosocin are identical between the vacant and terminating ribosome. Indeed, we observe that on the vacant 50S ribosome that the α-D-GalNAc modification has also inserted in between the U2609-A752 base-pair, causing a shift in U2609 as observed in the termination state (**Fig. 7b**). By contrast, the C-terminus of drosocin on the vacant 50S subunit appears flexible and less well-resolved, presumably, because the interaction with Gln235 of RF1 is absent (**Fig. 7b**). Similarly, binding of Api137 has previously been shown to be stabilized on 70S ribosomes by the presence of RF1 when compared to vacant ones (Florin et al., 2017). Since we observe no initiation states within our structural ensembles, we presume that the fMet-tRNA can bind at the P-site of the PTC unimpeded by the presence of Dro (**Fig. 7c**), presumably by effectively competing with the C-terminus of Dro for its binding site at the PTC. By contrast, we observe a major population of drosocin-bound ribosomes that are in an elongation state, namely, a post-peptide bond formation pre-translocation state with deacylated-tRNA^fMet^ in the P-site and a fMet-Leu-tRNA^Leu^ in the A-site (**Fig. 7d**). This suggests that drosocin interferes with the first translocation event that entails the movement of the fMet-Leu-tRNA^Leu^ into the P-site of the PTC. We believe that this arrest is likely to be temporary since in our toeprinting experiments, we observe that ribosomes can eventually translate the entire open reading frame and become trapped at the termination codon (**Fig. 1f-h**). In the elongation state, drosocin is particular flexible and poorly resolved, which is exemplified by the poor density for the α-D-GalNAc modification and the presence of both closed (base-paired) and open (unpaired) conformations of U2609 (**Fig. 7d** and **Supplementary Fig. 9**). We favour a model whereby drosocin and the fMet-Leu-tRNA^Leu^ jostle for position at the P-site of the PTC and that occupation by fMet-Leu-tRNA^Leu^ triggers translocation and subsequent rounds of elongation that ultimately cause dissociation of drosocin from the ribosome (**Fig. 7e**). Once the nascent polypeptide chain becomes extended within the ribosomal tunnel, drosocin cannot rebind until the termination codon is reached and the nascent chain is released by RF1 (or RF2) (**Fig. 7a**).

Collectively, our study demonstrates the advantage of forming complexes using less biased *in vitro* translation approaches, rather than vacant ribosomes or pre-defined termination states, enabling insights into early elongation events where the drosocin peptide competes with the growing nascent polypeptide chain. We believe that such events are likely to also exist for apidaecin since in addition to strong termination inhibition, some initiation effects have also been observed for apidaecin, both *in vivo* and *in vitro* (Mangano et al., 2020). It is remarkable that while both drosocin and apidaecin inhibit translation by trapping RFs on the ribosome in an analogous manner, the binding mode and molecular details of the interactions of these peptides with components of the ribosomal tunnel are completely distinct. This is accentuated by the presence of O-glycosylation that plays a critical role for drosocin, but is lacking for apidaecin. Curiously, there are other AMPs that are glycosylated, including diptericin, lebocin, formaecins and pyrrhocoricin (Bulet et al., 1999). In this regard, the PrAMP pyrrhocoricin is particularly relevant since it bears an identical modification to drosocin at exactly the same position, namely, GalNAc on Thr11, and minor forms with an additional galactose on the GalNAc have been also detected (Cociancich et al., 1994). Although structures of the unmodified pyrrhocoricin on the ribosome reveal a reversed orientation compared to drosocin (Gagnon et al., 2016; Seefeldt et al., 2016), superimposition reveals that Thr11 of pyrrhocoricin and drosocin are in close proximity, raising the possibility that the glycosylation of pyrrhocoricin may establish analogous interactions with the ribosome as observed here for drosocin. Lastly, we show that drosocin traps RF1 on the ribosome decoding the UAA stop codon in an analogous manner to that observed during canonical translation. However, the higher resolution observed here enables us to observe many water-mediated interactions that were not possible to observe previously. Thus, our study also provides structural insight into the fundamental mechanism of stop codon recognition during canonical translation termination.

## Methods

### Drosocin peptides

Api137, Dro, Dro-3P, Dro-R18A and Dro-R18K were synthesized by NovoPro (https://www.novoprolabs.com). The glycosylated Dro1-Dro8 peptides were synthesized as described (Lele et al., 2015a; Lele et al., 2017; Lele et al., 2015b; Talat et al., 2011)

### Bacterial strains

Strains *E. coli* Keio wt and *E. coli* Keio Δ*sbmA* used from Keio knockout collection (Horizon, a PerkinElmer Company; https://horizondiscovery.com). Wildtype *E. coli* SQ110 and SQ171 strains and related mutants *E. coli* SQ110 A2059G and *E. coli* SQ171 A2503G were obtained from the previous Api137 study (Florin et al., 2017). *E. coli* Strains SQ171 bearing A752G, U2609C and A752G:U2609C mutations (Svetlov et al., 2020) and *E. coli* SQ171 U2609G (Osterman et al., 2020) were generated previously.

### Antibiotic susceptibility assays

The susceptibility of *E. coli* strains to compounds was evaluated by monitoring the bacterial growth in presence of increasing concentrations of the compound of interest. Briefly, bacteria were inoculated in a total volume of 100 μL of medium contained in a well of a 96-well microplate (round bottom, with cap, sterile SARSTEDT). The medium used was either LB, as rich medium, or ATCC medium (778 Davis and Mingioli glucose minimal medium), as minimal medium. Before inoculation, bacteria were grown up to exponential phase and then inoculated into the culture mix, containing selective antibiotic if necessary, with an initial OD_600_ of 0.05. Values measured from wells containing just the medium were used as a blank. The growth in each well was monitored by measuring the OD_600_ every 10 mins for a total of 20 hours at 37 °C with shaking using a plate reader (Tecan Infinite®200 Pro). The inhibition resulting from a compound’s concentration was evaluated by normalizing the OD_600_ at t = 12 h (corresponding to the end of log phase) from the treated culture to the untreated one. For each compound, the concentration tested were 5 μM, 10 μM and 30 μM. Each single titration assay was done in triplicate with individually prepared culture mixes. For each concentration, the standard deviation was calculated taking into account each single replica and its specific technical error from the plate reader.

### Data analysis

Data from the *in vivo* assay were normalized and statistically analysed by GraphPad Prism V9.4.0.

### In vitro translation assays

The *in vitro* translation assay was carried out as described previously (Mardirossian et al., 2018b; Seefeldt et al., 2015) using the *E. coli* PURExpress®system (NEB E6800S). 1 µL of antibiotic solution was added to 5 µL of PURExpress® reaction mix. Each reaction contained 10 ng/μL of mRNA encoding the firefly luciferase, which was *in vitro* transcribed from a pIVEX-2.3MCS vector containing the firefly luciferase gene using T7 polymerase (ThermoScientific). The reaction mix was incubated for 30 min at 32 °C while shaking (600 rpm). Reactions were stopped with 5 µL kanamycin (50 mg/mL) and transferred into a 96-well microplate (Greiner Lumitrac, non-binding, white, chimney). 40 µL of luciferase assay substrate solution (Promega E1501) was added and luminescence was measured using a plate reader (Tecan Infinite®200 Pro). Nuclease free water was added instead of antibiotic as control. Absolute luminescence values were normalized using reactions without antibiotic. All assays were done as triplicates with individually prepared reaction mix.

### Toeprinting assays

Toeprinting reactions were performed as described previously (Seefeldt et al., 2015; Starosta et al., 2014). Briefly, reactions were performed with 6 μl of PURExpress ΔRF123 *in vitro* protein synthesis system (New England Biolabs) in the presence of 1x RF3, and either 1x, 2x or 10x of RF1 or RF2 (relative to the manufacturer’s recommendation). The reactions were carried out on MLIF-UAA-toeprint template (5’-TAATACGACTCACTATAGGGAGACTTAAGTATAAGGAGGAAAAAAT<u>**ATG**</u>ATATTCTTG<u>**T AAA**</u>TGCGTAATGTAGATAAAACATCTACTATTTAAGTGATAGAATTCTATCGTTAATAAGCAAAATTCATTATAACC-3’, ORF start- and stop-codon are underlined bold), containing T7 promotor, RBS, a MLIF coding ORF and the NV1* primer binding site. The template is a version of the ErmBL template previously described (Arenz et al., 2014) with a truncated ORF and addition of a isoleucine coding codon at the third position in the ORF. The template was generated by PCR of two overlapping 77 and 78 nt long primers. The reactions contained 30 ng of the MLIF-UAA-toeprint DNA template. The reactions were supplemented Api137, thiostrepton or one of the drosocin derivates as specified. The transcription-translation reactions were incubated for 15 min at 37°C. The reverse transcription on the MLIF-short-UAA toeprint template was carried out using AMV RT and primer NV*1-alexa647 (5’-GGTTATAATGAATTTTGCTTATTAAC-3’) previously described (Ramu et al., 2011). The transcription-translation reactions were incubated with AMV RT and NV*1-alexa647 for 20 min at 37 °C. mRNA degradation was carried out by addition of 1 µL of 5 M NaOH. The reactions were neutralized with 0.7 µL 25% HCl and nucleotide removal was performed with the QIAquick nucleotide removal kit (QIAGEN). The samples were dried under vacuum for 2 hours at 60°C for subsequent gel electrophoresis. The 6% acrylamide gels were scanned on a Typhoon scanner (GE Healthcare).

### Preparation of complexes for structural analysis

Drosocin-ribosome complexes were generated by *in vitro* transcription-translation reactions in PURExpress ΔRF123 *in vitro* protein synthesis system (New England Biolabs) with the same reaction mix as described earlier in the *toeprinting assays*. Complex formation reactions were carried out on MLIF-UAA toeprint DNA template in a 48 µL reaction with 1x RF3 and 10x RF1 (amounts relative to the manufacturer’s recommendation) in presence of 30 µM Dro1. The reaction was incubated for 15 min at 37°C. The reaction volume was then split: 42 µL were used for complex generation and 6 µL were used for toeprinting analysis. Ribosome complexes were isolated by centrifugation in 900 µL sucrose gradient buffer (containing 40% sucrose, 50 mM HEPES-KOH, pH 7.4, 100 mM KOAc, 25 mM Mg(OAc)_2_ and 6 mM 2-mercaptoethanol) for 3 hours at 4°C with 80,000 x g in a Optima™ Max-XP Tabletop Ultracentrifuge with a TLA 120.2 rotor. The pelleted complex was resuspended in Hico buffer (50 mM HEPES-KOH, pH 7.4, 100 mM KOAc, 25 mM Mg(OAc)_2_ supplemented with RF1, RF3 and Dro1 at the same concentrations used in the *in vitro* translation reaction), then incubated for 15 min at 37°C.

### Preparation of cryo-EM grids and data collection

Grids (Quantifoil R3/3 Cu300 with 3 nm holey carbon) were glow discharged and 4 µL of sample (8 OD_260_/mL) was applied using a Vitrobot Mark IV (FEI) and snap frozen in ethane/propane. Frozen cryo-EM grids were imaged on a TFS 300kV Titan Krios at the Dubochet Center for Imaging EPFL (Lausanne, Switzerland). Images were collected on Falcon IV direct detection camera in counting mode using the EPU and AFIS data collection scheme with a magnification of 96,000 x and a total dose of 40 electrons per square angstrom (e^-^/Å^2^) for each exposure, and defocus ranging from -0.4 to -0.9 microns. In total, 8,861 movies were produced in EER format.

### Single-particle reconstruction of drosocin-ribosome complexes

RELION v4.0 (Kimanius et al., 2021; Zivanov et al., 2018) was used for processing, unless otherwise specified. For motion correction, RELION’s implementation of MotionCor2 with 4×4 patches and for initial CTF estimation CTFFIND v4.1.14 (Rohou and Grigorieff, 2015; Zheng et al., 2017) was employed. From 8,861 micrographs, 715,455 particles were picked using crYOLO with a general model (Wagner et al., 2019). 529,600 ribosome-like particles were selected after 2D classification and extracted at 3x decimated pixel size (2.4 Å/pixel). An initial 3D refinement was done using a *E. coli* 70S reference map (EMD-12573) (Beckert et al., 2021) and followed by initial 3D classification without angular sampling with six classes. Two classes containing 70S ribosomes were combined (356,671 particles) and sub-sorted. A class containing 50S subunits (159,749 particles) was further processed. We observed no classes containing RF3, despite the presence of RF3 in the translation reactions. However, unlike our previous study (Graf et al., 2018), we did not use non-hydrolysable GTP analogs. The sub-sorting was done using particle subtraction with a circular mask around the A-site with four classes. Classes containing density that could be assigned RF1 (137,449 particles) and one class with A-tRNA density (84,697 particles) were further processed. All resulting classes were 3D refined and CTF refined (4^th^ order aberrations, beam-tilt, anisotropic magnification and per-particle defocus value estimation). The termination complex was additionally subjected to Bayesian polishing (Zivanov et al., 2019) and another round of CTF refinement. For the termination, elongation and 50S complexes final resolutions (Gold-standard FSC_0.143_) of masked reconstructions of 2.3 Å, 2.8 Å and 2.1 Å were achieved respectively. To estimate local resolution values Bsoft (Heymann, 2018) was used on the half-maps of the final reconstructions (blocres -sampling 0.8 -maxres -boc 20 -cutoff 0.143 -verbose 1 -origin 0,0,0 -Mask half_map1 half_map 2).

### Molecular modelling of the drosocin-ribosome complexes

The molecular models of the 30S and 50S ribosomal subunits were based on a high resolution *E. coli* 70S ribosome (PDB ID 7K00) (Watson et al., 2020). Drosocin was modelled *de novo* and the 2-acetamido-2-deoxy-alpha-D-galactopyranose was taken from the ligand expo database A2G (PDB ID 1D0H) (Emsley et al., 2000) and linked through REFMAC 5 (Vagin et al., 2004). Restraints files for modified residues were created using aceDRG (Long et al., 2017). The termination complex was assembled with a RF1 Alphafold model (AF-P0A7I0-F1)(Jumper et al., 2021; Varadi et al., 2022) and a crystal structure of a deacylated phenylalanine tRNA (PDB ID 6Y3G)(Bourgeois et al., 2020) in the P-site. The elongation complex was assembled with an initiator fMet-tRNA (PDB ID 1VY4) (Polikanov et al., 2014) in the P-site and a Leu-tRNA (PDB ID 7NSQ) (Beckert et al., 2021) in the A-site. Starting models were rigid body fitted using ChimeraX (Goddard et al., 2018; Pettersen et al., 2021) and modelled using Coot 0.9.8.3 (Emsley et al., 2010) from the CCP4 software suite v.8.0 (Winn et al., 2011). The sequence for the tRNAs were adjusted based the appropriate anticodons corresponding to the mRNA. Final refinements were done in REFMAC 5 (Vagin et al., 2004) using Servalcat (Yamashita et al., 2021). The molecular models were validated using Phenix comprehensive Cryo-EM validation in Phenix 1.20-4487 (Chen et al., 2010; Liebschner et al., 2019).

### Figures

UCSF ChimeraX 1.3 (Goddard et al., 2018) was used to isolate density and visualize density images and structural superpositions. Models were aligned using PyMol v2.4 (Schrödinger, LLC). Figures were assembled with Adobe Illustrator (Adobe Inc.) and Inkscape (latest development release, regularly updated).

## Data availability

Micrographs have been deposited as uncorrected frames in the Electron Microscopy Public Image Archive (EMPIAR) with the accession codes EMPIAR-XXXXX [https://www.ebi.ac.uk/pdbe/emdb/empiar/entry/10764/]. Cryo-EM maps have been deposited in the Electron Microscopy Data Bank (EMDB) with accession codes EMD-XXXX [https://www.ebi.ac.uk/pdbe/entry/emdb/EMD-1XXX] (Drosocin-termination complex), EMD-YYYY [https://www.ebi.ac.uk/pdbe/entry/emdb/EMD-13242] (Drosocin-elongation complex), and EMD-ZZZZ [https://www.ebi.ac.uk/pdbe/entry/emdb/EMD-13243] (Drosocin-50S complex). Molecular models have been deposited in the Protein Data Bank with accession codes 8XYZ [https://doi.org/10.2210/pdb8A57/pdb] (Drosocin-termination complex), 8XYZ [https://doi.org/10.2210/pdb8A63/pdb] (Drosocin-elongation complex), 8XYZ [https://doi.org/10.2210/pdb8A5I/pdb] (Drosocin-50S complex).

Source data are provided with this paper.

## Acknowledgments

We thank A. Myasnikov, S. Nazarov and E. Ushikawa from the Dubochet Center for Imaging (an EPFL, UNIGE, UNIL initiative in Lausanne, Switzerland) for help with cryo-EM data collection, data processing and IT support. D.N.W. is supported by the Deutsche Zentrum für Luft-und Raumfahrt (DLR01Kl1820) within the RIBOTARGET consortium under the framework of JPIAMR.

## Author contributions

D.S.L. and K.J.K. synthesized modified drosocin peptides. M.M. performed all growth assays and *in vitro* translation assays. M.B. prepared cryo-EM sample and performed toeprinting analysis. H.S. prepared cryo-EM grids and B.B. collected the cryo-EM data. T.O.K. processed the microscopy data, generated and refined the molecular models. T.O.K, M.M. and M.B. prepared the figures. D.N.W. wrote the manuscript with input from all authors. D.N.W conceived and supervised the project.

## Additional information

**Supplementary information** The online version contains supplementary material available at https://…

**Correspondence** and requests for materials should be addressed to D.N.W.

## References

Ahn, M., Murugan, R.N., Nan, Y.H., Cheong, C., Sohn, H., Kim, E.H., Hwang, E., Ryu, E.K., Kang, S.W., Shin, S.Y., et al. (2011a). Substitution of the GalNAc-alpha-O-Thr(1)(1) residue in drosocin with O-linked glyco-peptoid residue: effect on antibacterial activity and conformational change. Bioorg Med Chem Lett 21, 6148–6153.

Ahn, M., Sohn, H., Nan, Y.H., Murugan, R.N., Cheong, C., Eun Kyoung Ryu, Kim, E.-H., Kang, S.W., Kim, E.J., Shin, S.Y., et al. (2011b). Functional and Structural Characterization of Drosocin and its Derivatives Linked O-GalNAc at Thr11 Residue. Bull. Korean Chem. Soc. 32, 3327–3332.

Arenz, S., Ramu, H., Gupta, P., Berninghausen, O., Beckmann, R., Vazquez-Laslop, N., Mankin, A.S., and Wilson, D.N. (2014). Molecular basis for erythromycin-dependent ribosome stalling during translation of the ErmBL leader peptide. Nat Commun 5, 3501.

Beckert, B., Leroy, E.C., Sothiselvam, S., Bock, L.V., Svetlov, M.S., Graf, M., Arenz, S., Abdelshahid, M., Seip, B., Grubmuller, H., et al. (2021). Structural and mechanistic basis for translation inhibition by macrolide and ketolide antibiotics. Nat Commun 12, 4466.

Benincasa, M., Scocchi, M., Podda, E., Skerlavaj, B., Dolzani, L., and Gennaro, R. (2004). Antimicrobial activity of Bac7 fragments against drug-resistant clinical isolates. Peptides 25, 2055–2061.

Berthold, N., and Hoffmann, R. (2014). Cellular uptake of apidaecin 1b and related analogs in Gram-negative bacteria reveals novel antibacterial mechanism for proline-rich antimicrobial peptides. Protein Pept Lett 21, 391–398.

Bikker, F.J., Kaman-van Zanten, W.E., de Vries-van de Ruit, A.M., Voskamp-Visser, I., van Hooft, P.A., Mars-Groenendijk, R.H., de Visser, P.C., and Noort, D. (2006). Evaluation of the antibacterial spectrum of drosocin analogues. Chem Biol Drug Des 68, 148–153.

Bourgeois, G., Seguin, J., Babin, M., Gondry, M., Mechulam, Y., and Schmitt, E. (2020). Structural basis of the interaction between cyclodipeptide synthases and aminoacylated tRNA substrates. RNA 26, 1589–1602.

Bulet, P., Dimarcq, J.L., Hetru, C., Lagueux, M., Charlet, M., Hegy, G., Van Dorsselaer, A., and Hoffmann, J.A. (1993). A novel inducible antibacterial peptide of Drosophila carries an O-glycosylated substitution. J Biol Chem 268, 14893–14897.

Bulet, P., Hetru, C., Dimarcq, J.L., and Hoffmann, D. (1999). Antimicrobial peptides in insects; structure and function. Dev Comp Immunol 23, 329–344.

Bulet, P., Urge, L., Ohresser, S., Hetru, C., and Otvos, L., Jr. (1996). Enlarged scale chemical synthesis and range of activity of drosocin, an O-glycosylated antibacterial peptide of Drosophila. Eur J Biochem 238, 64–69.

Castle, M., Nazarian, A., Yi, S.S., and Tempst, P. (1999). Lethal effects of apidaecin on Escherichia coli involve sequential molecular interactions with diverse targets. J Biol Chem 274, 32555–32564.

Chan, K.H., Petrychenko, V., Mueller, C., Maracci, C., Holtkamp, W., Wilson, D.N., Fischer, N., and Rodnina, M.V. (2020). Mechanism of ribosome rescue by alternative ribosome-rescue factor B. Nat Commun 11, 4106.

Chen, V.B., Arendall, W.B., 3rd, Headd, J.J., Keedy, D.A., Immormino, R.M., Kapral, G.J., Murray, L.W., Richardson, J.S., and Richardson, D.C. (2010). MolProbity: all-atom structure validation for macromolecular crystallography. Acta Crystallogr D Biol Crystallogr 66, 12–21.

Cociancich, S., Dupont, A., Hegy, G., Lanot, R., Holder, F., Hetru, C., Hoffmann, J.A., and Bulet, P. (1994). Novel inducible antibacterial peptides from a hemipteran insect, the sap-sucking bug Pyrrhocoris apterus. Biochem J 300 (Pt 2), 567–575.

de Visser, P.C., van Hooft, P.A., de Vries, A.M., de Jong, A., van der Marel, G.A., Overkleeft, H.S., and Noort, D. (2005). Biological evaluation of Tyr6 and Ser7 modified drosocin analogues. Bioorg Med Chem Lett 15, 2902–2905.

Dunkle, J.A., Xiong, L., Mankin, A.S., and Cate, J.H. (2010). Structures of the Escherichia coli ribosome with antibiotics bound near the peptidyl transferase center explain spectra of drug action. Proc Natl Acad Sci U S A 107, 17152–17157.

Emsley, P., Fotinou, C., Black, I., Fairweather, N.F., Charles, I.G., Watts, C., Hewitt, E., and Isaacs, N.W. (2000). The structures of the H(C) fragment of tetanus toxin with carbohydrate subunit complexes provide insight into ganglioside binding. J Biol Chem 275, 8889–8894.

Emsley, P., Lohkamp, B., Scott, W.G., and Cowtan, K. (2010). Features and development of Coot. Acta Crystallogr D Biol Crystallogr 66, 486–501.

Florin, T., Maracci, C., Graf, M., Karki, P., Klepacki, D., Berninghausen, O., Beckmann, R., Vazquez-Laslop, N., Wilson, D.N., Rodnina, M.V., et al. (2017). An antimicrobial peptide that inhibits translation by trapping release factors on the ribosome. Nat Struct Mol Biol 24, 752–757.

Fu, Z., Indrisiunaite, G., Kaledhonkar, S., Shah, B., Sun, M., Chen, B., Grassucci, R.A., Ehrenberg, M., and Frank, J. (2019). The structural basis for release-factor activation during translation termination revealed by time-resolved cryogenic electron microscopy. Nat Commun 10, 2579.

Gagnon, M.G., Roy, R.N., Lomakin, I.B., Florin, T., Mankin, A.S., and Steitz, T.A. (2016). Structures of proline-rich peptides bound to the ribosome reveal a common mechanism of protein synthesis inhibition. Nucleic acids research 44, 2439–2450.

Gobbo, M., Biondi, L., Filira, F., Gennaro, R., Benincasa, M., Scolaro, B., and Rocchi, R. (2002). Antimicrobial peptides: synthesis and antibacterial activity of linear and cyclic drosocin and apidaecin 1b analogues. J Med Chem 45, 4494–4504.

Goddard, T.D., Huang, C.C., Meng, E.C., Pettersen, E.F., Couch, G.S., Morris, J.H., and Ferrin, T.E. (2018). UCSF ChimeraX: Meeting modern challenges in visualization and analysis. Protein Sci 27, 14–25.

Graf, M., Huter, P., Maracci, C., Peterek, M., Rodnina, M.V., and Wilson, D.N. (2018). Visualization of translation termination intermediates trapped by the Apidaecin 137 peptide during RF3-mediated recycling of RF1. Nat Commun 9, 3053.

Graf, M., Mardirossian, M., Nguyen, F., Seefeldt, A.C., Guichard, G., Scocchi, M., Innis, C.A., and Wilson, D.N. (2017). Proline-rich antimicrobial peptides targeting protein synthesis. Natural product reports 34, 702–711.

Graf, M., and Wilson, D.N. (2019). Intracellular Antimicrobial Peptides Targeting the Protein Synthesis Machinery. Adv Exp Med Biol 1117, 73–89.

Hartz, D., McPheeters, D.S., Traut, R., and Gold, L. (1988). Extension inhibition analysis of translation initiation complexes. Methods Enzymol. 164, 419–425.

Heymann, J.B. (2018). Guidelines for using Bsoft for high resolution reconstruction and validation of biomolecular structures from electron micrographs. Protein Sci 27, 159–171.

Hoffmann, R., Bulet, P., Urge, L., and Otvos, L., Jr. (1999). Range of activity and metabolic stability of synthetic antibacterial glycopeptides from insects. Biochim Biophys Acta 1426, 459–467.

Jumper, J., Evans, R., Pritzel, A., Green, T., Figurnov, M., Ronneberger, O., Tunyasuvunakool, K., Bates, R., Zidek, A., Potapenko, A., et al. (2021). Highly accurate protein structure prediction with AlphaFold. Nature 596, 583–589.

Kimanius, D., Dong, L., Sharov, G., Nakane, T., and Scheres, S.H.W. (2021). New tools for automated cryo-EM single-particle analysis in RELION-4.0. Biochem J 478, 4169–4185.

Krizsan, A., Prahl, C., Goldbach, T., Knappe, D., and Hoffmann, R. (2015). Short Proline-Rich Antimicrobial Peptides Inhibit Either the Bacterial 70S Ribosome or the Assembly of its Large 50S Subunit. Chembiochem 16, 2304–2308.

Krizsan, A., Volke, D., Weinert, S., Strater, N., Knappe, D., and Hoffmann, R. (2014). Insect-derived proline-rich antimicrobial peptides kill bacteria by inhibiting bacterial protein translation at the 70S ribosome. Angew Chem Int Ed Engl 53, 12236–12239.

Laurberg, M., Asahara, H., Korostelev, A., Zhu, J., Trakhanov, S., and Noller, H.F. (2008). Structural basis for translation termination on the 70S ribosome. Nature 454, 852–857.

Lele, D.S., Dwivedi, R., Kumari, S., and Kaur, K.J. (2015a). Effect of distal sugar and interglycosidic linkage of disaccharides on the activity of proline rich antimicrobial glycopeptides. J Pept Sci 21, 833–844.

Lele, D.S., Kaur, G., Thiruvikraman, M., and Kaur, K.J. (2017). Comparing naturally occurring glycosylated forms of proline rich antibacterial peptide, Drosocin. Glycoconj J 34, 613–624.

Lele, D.S., Talat, S., and Kaur, K.J. (2013). The Presence of Arginine in the Pro-Arg-Pro Motif Augments the Lethality of Proline Rich Antimicrobial Peptides of Insect Source. Int. J. Pept. Res. Ther. 19, 323–330.

Lele, D.S., Talat, S., Kumari, S., Srivastava, N., and Kaur, K.J. (2015b). Understanding the importance of glycosylated threonine and stereospecific action of Drosocin, a Proline rich antimicrobial peptide. European journal of medicinal chemistry 92, 637–647.

Liebschner, D., Afonine, P.V., Baker, M.L., Bunkoczi, G., Chen, V.B., Croll, T.I., Hintze, B., Hung, L.W., Jain, S., McCoy, A.J., et al. (2019). Macromolecular structure determination using X-rays, neutrons and electrons: recent developments in Phenix. Acta Crystallogr D Struct Biol 75, 861–877.

Long, F., Nicholls, R.A., Emsley, P., Graaeulis, S., Merkys, A., Vaitkus, A., and Murshudov, G.N. (2017). AceDRG: a stereochemical description generator for ligands. Acta Crystallogr D Struct Biol 73, 112–122.

Ludwig, T., Krizsan, A., Mohammed, G.K., and Hoffmann, R. (2022). Antimicrobial Activity and 70S Ribosome Binding of Apidaecin-Derived Api805 with Increased Bacterial Uptake Rate. Antibiotics (Basel) 11.

Mangano, K., Florin, T., Shao, X., Klepacki, D., Chelysheva, I., Ignatova, Z., Gao, Y., Mankin, A.S., and Vazquez-Laslop, N. (2020). Genome-wide effects of the antimicrobial peptide apidaecin on translation termination in bacteria. eLife 9.

Marcaurelle, L.A., E.C., R., and Bertozzi, C. (1998). Synthesis of an oxime-linked neoglycopeptide with glycosylation-dependent activity similar to its native counterpart. Tetrahedron Letters 39, 8417–8420.

Mardirossian, M., Barriere, Q., Timchenko, T., Muller, C., Pacor, S., Mergaert, P., Scocchi, M., and Wilson, D.N. (2018a). Fragments of the Nonlytic Proline-Rich Antimicrobial Peptide Bac5 Kill Escherichia coli Cells by Inhibiting Protein Synthesis. Antimicrob Agents Chemother 62.

Mardirossian, M., Grzela, R., Giglione, C., Meinnel, T., Gennaro, R., Mergaert, P., and Scocchi, M. (2014). The host antimicrobial peptide Bac71-35 binds to bacterial ribosomal proteins and inhibits protein synthesis. Chem Biol 21, 1639–1647.

Mardirossian, M., Perebaskine, N., Benincasa, M., Gambato, S., Hofmann, S., Huter, P., Muller, C., Hilpert, K., Innis, C.A., Tossi, A., et al. (2018b). The Dolphin Proline-Rich Antimicrobial Peptide Tur1A Inhibits Protein Synthesis by Targeting the Bacterial Ribosome. Cell Chem Biol 25, 530–539 e537.

Mardirossian, M., Sola, R., Beckert, B., Collis, D.W.P., Di Stasi, A., Armas, F., Hilpert, K., Wilson, D.N., and Scocchi, M. (2019). Proline-Rich Peptides with Improved Antimicrobial Activity against E. coli, K. pneumoniae, and A. baumannii. ChemMedChem 14, 2025–2033.

Mardirossian, M., Sola, R., Beckert, B., Valencic, E., Collis, D.W.P., Borisek, J., Armas, F., Di Stasi, A., Buchmann, J., Syroegin, E.A., et al. (2020). Peptide Inhibitors of Bacterial Protein Synthesis with Broad Spectrum and SbmA-Independent Bactericidal Activity against Clinical Pathogens. J Med Chem 63, 9590–9602.

Mattiuzzo, M., Bandiera, A., Gennaro, R., Benincasa, M., Pacor, S., Antcheva, N., and Scocchi, M. (2007). Role of the Escherichia coli SbmA in the antimicrobial activity of proline-rich peptides. Mol Microbiol 66, 151–163.

McManus, A.M., Otvos, L., Jr., Hoffmann, R., and Craik, D.J. (1999). Conformational studies by NMR of the antimicrobial peptide, drosocin, and its non-glycosylated derivative: effects of glycosylation on solution conformation. Biochemistry 38, 705–714.

Osterman, I.A., Wieland, M., Maviza, T.P., Lashkevich, K.A., Lukianov, D.A., Komarova, E.S., Zakalyukina, Y.V., Buschauer, R., Shiriaev, D.I., Leyn, S.A., et al. (2020). Tetracenomycin X inhibits translation by binding within the ribosomal exit tunnel. Nat Chem Biol 16, 1071–1077.

Otvos, L., Jr., O, I., Rogers, M.E., Consolvo, P.J., Condie, B.A., Lovas, S., Bulet, P., and Blaszczyk-Thurin, M. (2000). Interaction between heat shock proteins and antimicrobial peptides. Biochemistry 39, 14150–14159.

Pettersen, E.F., Goddard, T.D., Huang, C.C., Meng, E.C., Couch, G.S., Croll, T.I., Morris, J.H., and Ferrin, T.E. (2021). UCSF ChimeraX: Structure visualization for researchers, educators, and developers. Protein Sci 30, 70–82.

Pierson, W.E., Hoffer, E.D., Keedy, H.E., Simms, C.L., Dunham, C.M., and Zaher, H.S. (2016). Uniformity of Peptide Release Is Maintained by Methylation of Release Factors. Cell reports 17, 11–18.

Polikanov, Y.S., Steitz, T.A., and Innis, C.A. (2014). A proton wire to couple aminoacyl-tRNA accommodation and peptide-bond formation on the ribosome. Nat Struct Mol Biol 21, 787–793.

Rabel, D., Charlet, M., Ehret-Sabatier, L., Cavicchioli, L., Cudic, M., Otvos, L., Jr., and Bulet, P. (2004). Primary structure and in vitro antibacterial properties of the Drosophila melanogaster attacin C Pro-domain. J Biol Chem 279, 14853–14859.

Ramu, H., Vazquez-Laslop, N., Klepacki, D., Dai, Q., Piccirilli, J., Micura, R., and Mankin, A.S. (2011). Nascent peptide in the ribosome exit tunnel affects functional properties of the A-site of the peptidyl transferase center. Mol Cell 41, 321–330.

Rodriguez, E.C., Winans, K.A., King, D.S., and Bertozzi, C.R. (1997). A Strategy for the Chemoselective Synthesis of O-Linked Glycopeptides with Native Sugar−Peptide Linkages. J. Am. Chem. Soc. 113, 9905–9906.

Rohou, A., and Grigorieff, N. (2015). CTFFIND4: Fast and accurate defocus estimation from electron micrographs. J Struct Biol 192, 216–221.

Roy, R.N., Lomakin, I.B., Gagnon, M.G., and Steitz, T.A. (2015). The mechanism of inhibition of protein synthesis by the proline-rich peptide oncocin. Nat Struct Mol Biol 22, 466–469.

Runti, G., Lopez Ruiz Mdel, C., Stoilova, T., Hussain, R., Jennions, M., Choudhury, H.G., Benincasa, M., Gennaro, R., Beis, K., and Scocchi, M. (2013). Functional characterization of SbmA, a bacterial inner membrane transporter required for importing the antimicrobial peptide Bac7(1-35). J Bacteriol 195, 5343–5351.

Scocchi, M., Tossi, A., and Gennaro, R. (2011). Proline-rich antimicrobial peptides: converging to a non-lytic mechanism of action. Cell Mol Life Sci 68, 2317–2330.

Seefeldt, A.C., Graf, M., Perebaskine, N., Nguyen, F., Arenz, S., Mardirossian, M., Scocchi, M., Wilson, D.N., and Innis, C.A. (2016). Structure of the mammalian antimicrobial peptide Bac7(1-16) bound within the exit tunnel of a bacterial ribosome. Nucleic acids research 44, 2429–2438.

Seefeldt, A.C., Nguyen, F., Antunes, S., Perebaskine, N., Graf, M., Arenz, S., Inampudi, K.K., Douat, C., Guichard, G., Wilson, D.N., et al. (2015). The proline-rich antimicrobial peptide Onc112 inhibits translation by blocking and destabilizing the initiation complex. Nat Struct Mol Biol 22, 470–475.

Sola, R., Mardirossian, M., Beckert, B., Sanghez De Luna, L., Prickett, D., Tossi, A., Wilson, D.N., and Scocchi, M. (2020). Characterization of Cetacean Proline-Rich Antimicrobial Peptides Displaying Activity against ESKAPE Pathogens. Int J Mol Sci 21.

Starosta, A.L., Lassak, J., Peil, L., Atkinson, G.C., Virumae, K., Tenson, T., Remme, J., Jung, K., and Wilson, D.N. (2014). Translational stalling at polyproline stretches is modulated by the sequence context upstream of the stall site. Nucleic Acids Res 42, 10711–10719.

Svetlov, M.S., Cohen, S., Alsuhebany, N., Vazquez-Laslop, N., and Mankin, A.S. (2020). A long-distance rRNA base pair impacts the ability of macrolide antibiotics to kill bacteria. Proc Natl Acad Sci U S A 117, 1971–1975.

Syroegin, E.A., Aleksandrova, E.V., and Polikanov, Y.S. (2022a). Structural basis for the inability of chloramphenicol to inhibit peptide bond formation in the presence of A-site glycine. Nucleic Acids Res 50, 7669–7679.

Syroegin, E.A., Flemmich, L., Klepacki, D., Vazquez-Laslop, N., Micura, R., and Polikanov, Y.S. (2022b). Structural basis for the context-specific action of the classic peptidyl transferase inhibitor chloramphenicol. Nat Struct Mol Biol 29, 152–161.

Talat, S., Thiruvikraman, M., Kumari, S., and Kaur, K.J. (2011). Glycosylated analogs of formaecin I and drosocin exhibit differential pattern of antibacterial activity. Glycoconj J 28, 537–555.

Uttenweiler-Joseph, S., Moniatte, M., Lagueux, M., Van Dorsselaer, A., Hoffmann, J.A., and Bulet, P. (1998). Differential display of peptides induced during the immune response of Drosophila: a matrix-assisted laser desorption ionization time-of-flight mass spectrometry study. Proc Natl Acad Sci U S A 95, 11342–11347.

Vagin, A.A., Steiner, R.A., Lebedev, A.A., Potterton, L., McNicholas, S., Long, F., and Murshudov, G.N. (2004). REFMAC5 dictionary: organization of prior chemical knowledge and guidelines for its use. Acta Crystallogr D Biol Crystallogr 60, 2184–2195.

Varadi, M., Anyango, S., Deshpande, M., Nair, S., Natassia, C., Yordanova, G., Yuan, D., Stroe, O., Wood, G., Laydon, A., et al. (2022). AlphaFold Protein Structure Database: massively expanding the structural coverage of protein-sequence space with high-accuracy models. Nucleic Acids Res 50, D439–D444.

Wagner, T., Merino, F., Stabrin, M., Moriya, T., Antoni, C., Apelbaum, A., Hagel, P., Sitsel, O., Raisch, T., Prumbaum, D., et al. (2019). SPHIRE-crYOLO is a fast and accurate fully automated particle picker for cryo-EM. Commun Biol 2, 218.

Watson, Z.L., Ward, F.R., Meheust, R., Ad, O., Schepartz, A., Banfield, J.F., and Cate, J.H. (2020). Structure of the bacterial ribosome at 2 A resolution. eLife 9.

Winn, M.D., Ballard, C.C., Cowtan, K.D., Dodson, E.J., Emsley, P., Evans, P.R., Keegan, R.M., Krissinel, E.B., Leslie, A.G., McCoy, A., et al. (2011). Overview of the CCP4 suite and current developments. Acta Crystallogr D Biol Crystallogr 67, 235–242.

Yamashita, K., Palmer, C.M., Burnley, T., and Murshudov, G.N. (2021). Cryo-EM single-particle structure refinement and map calculation using Servalcat. Acta Crystallogr D Struct Biol 77, 1282–1291.

Yang, K., Chang, J.Y., Cui, Z., Li, X., Meng, R., Duan, L., Thongchol, J., Jakana, J., Huwe, C.M., Sacchettini, J.C., et al. (2017). Structural insights into species-specific features of the ribosome from the human pathogen Mycobacterium tuberculosis. Nucleic Acids Res 45, 10884–10894.

Zheng, S.Q., Palovcak, E., Armache, J.P., Verba, K.A., Cheng, Y., and Agard, D.A. (2017). MotionCor2: anisotropic correction of beam-induced motion for improved cryo-electron microscopy. Nat Methods 14, 331–332.

Zhou, J., Korostelev, A., Lancaster, L., and Noller, H.F. (2012). Crystal structures of 70S ribosomes bound to release factors RF1, RF2 and RF3. Curr Opin Struct Biol 22, 733–742.

Zivanov, J., Nakane, T., Forsberg, B.O., Kimanius, D., Hagen, W.J., Lindahl, E., and Scheres, S.H. (2018). New tools for automated high-resolution cryo-EM structure determination in RELION-3. eLife 7.

Zivanov, J., Nakane, T., and Scheres, S.H.W. (2019). A Bayesian approach to beam-induced motion correction in cryo-EM single-particle analysis. IUCrJ 6, 5–17.

