## Supplementary Information for "Structural basis for translation inhibition by the glycosylated antimicrobial peptide Drosocin from *Drosophila melanogaster*"

**Table S1. Cryo-EM data collection, modelling and refinement statistics.**

| | Termination complex | Elongation complex | 50S complex |
|---|---|---|---|
| EMDB ID | EMD-15488 | EMD-15523 | EMD-15533 |
| PDB ID | 8AKN | 8AM9 | 8ANA |
| **Data collection** | | | |
| Magnification (×) | 96,000 | 96,000 | 96,000 |
| Electron fluence ($e^-/Å^2$) | 40 | 40 | 40 |
| Defocus range (µm) | -0.4 to -0.9 | -0.4 to -0.9 | -0.4 to -0.9 |
| Pixel size (Å) | 0.80 | 0.80 | 0.80 |
| Initial particles | 529,600 | 529,600 | 529,600 |
| Final particles | 137,449 | 84,697 | 159,749 |
| Average resolution (Å) (FSC threshold 0.143) | 2.3 | 2.8 | 2.1 |
| **Model composition** | | | |
| Atoms | 146,523 | 142,038 | 86,889 |
| Protein residues | 5,960 | 5,606 | 3,196 |
| RNA bases | 4,552 | 4,554 | 2,872 |
| **Refinement** | | | |
| Map CC around atoms | 0.69 | 0.82 | 0.84 |
| Map CC whole unit cell | 0.65 | 0.70 | 0.77 |
| Map sharpening B factor ($Å^2$) | -4.7 | -26.7 | -7.2 |
| R.M.S. deviations | | | |
| Bond lengths (Å) | 0.015 | 0.009 | 0.011 |
| Bond angles (°) | 1.863 | 1.014 | 1.659 |
| **Validation** | | | |
| MolProbity score | 1.88 | 2.00 | 1.40 |
| Clash score | 3.77 | 5.60 | 2.12 |
| Poor rotamers (%) | 2.15 | 3.55 | 1.23 |
| Ramachandran statistics | | | |
| Favoured (%) | 92.76 | 95.91 | 94.95 |
| Outlier (%) | 1.06 | 0.27 | 0.8 |

### Supplementary Figures

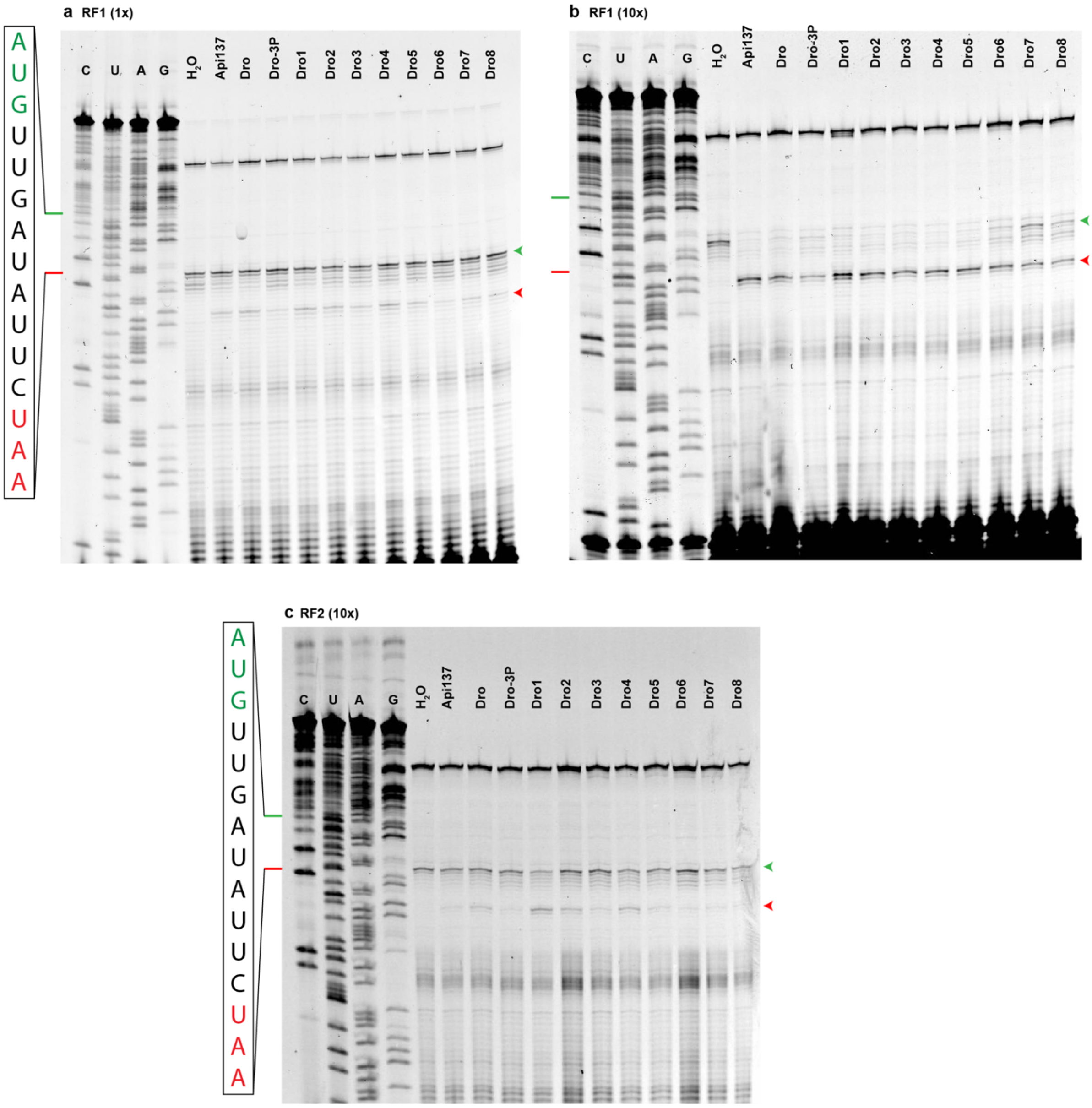


**Supplementary Fig. 1: Toeprint of drosocin in presence of release factors. a-c**, Uncropped toeprinting assays from figure 1g-h monitoring the position of ribosomes on an MLIF*-mRNA in the presence of 30 µM Api137 and drosocin derivatives and either (**a**) 1x RF1, (**b**) 10x RF1 or (**c**) 10x RF2. Bands corresponding to ribosomes present at the start and stop codons are indicated by green and red arrows, respectively. The start and stop codons on the sequencing lanes are indicated by green and red lines and correspond to the sequence shown on the left.

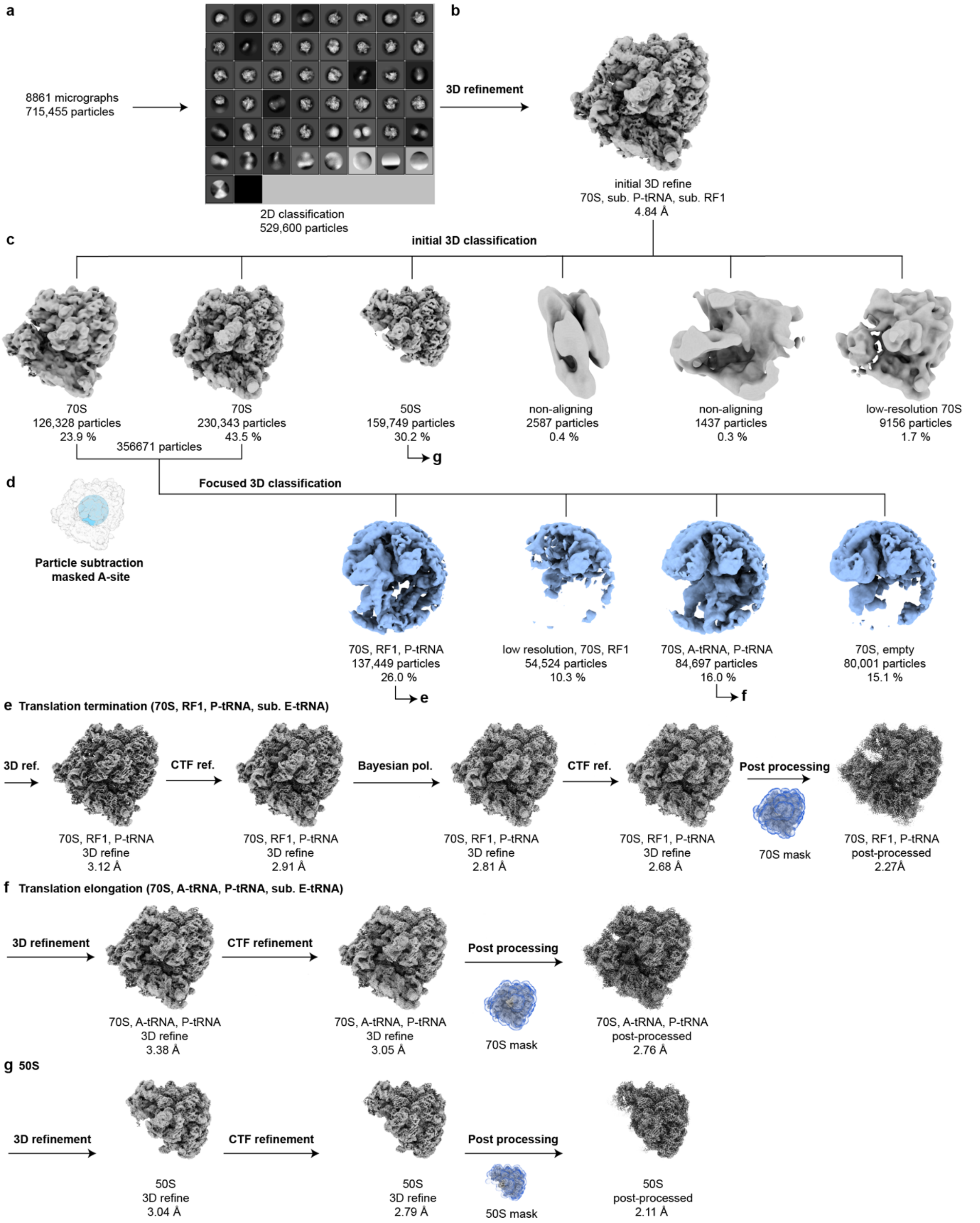


**Supplementary Fig. 2: *In silico* sorting scheme. a**, From 8,861 micrographs 715,455 particles were picked and subjected to 2D classification resulting in 529,600 ribosome-like particles. **b**, Particles were initially 3D refined at 3x decimated pixel size. **c**, Initial 3D classification for 70 iterations resulted in six classes. Classes containing 70S were combined and subsorted (356,671 particles) and a class containing 50S (159,749 particles) was further processed. **d**, 70S particles were subtracted with a mask around the A-site and subjected to 150 iterations of focused 3D classification. **e**, Termination complex containing RF1, P-tRNA and sub. E-tRNA density was 3D refined at undecimated pixel size and subjected to CTF refinement ($4^{th}$ order aberrations, beam-tilt, anisotropic magnification and per-particle defocus value estimation), Bayesian polished and again CTF refined resulting in a final average resolution for the masked reconstruction of 2.3 Å (at $FSC_{0.143}$). **f**, Elongation complex containing A-tRNA and P-tRNA density was 3D refined at undecimated pixel size and subjected to CTF refinement ($4^{th}$ order aberrations, beam-tilt, anisotropic magnification and per-particle defocus value estimation), resulting in a final average resolution for the masked reconstruction of 2.8 Å (at $FSC_{0.143}$). **g**, 50S complex containing A-tRNA and P-tRNA density was 3D refined at undecimated pixel size and subjected to CTF refinement ($4^{th}$ order aberrations, beam-tilt, anisotropic magnification and per-particle defocus value estimation), resulting in a final average resolution for the masked reconstruction of 2.1 Å (at $FSC_{0.143}$).

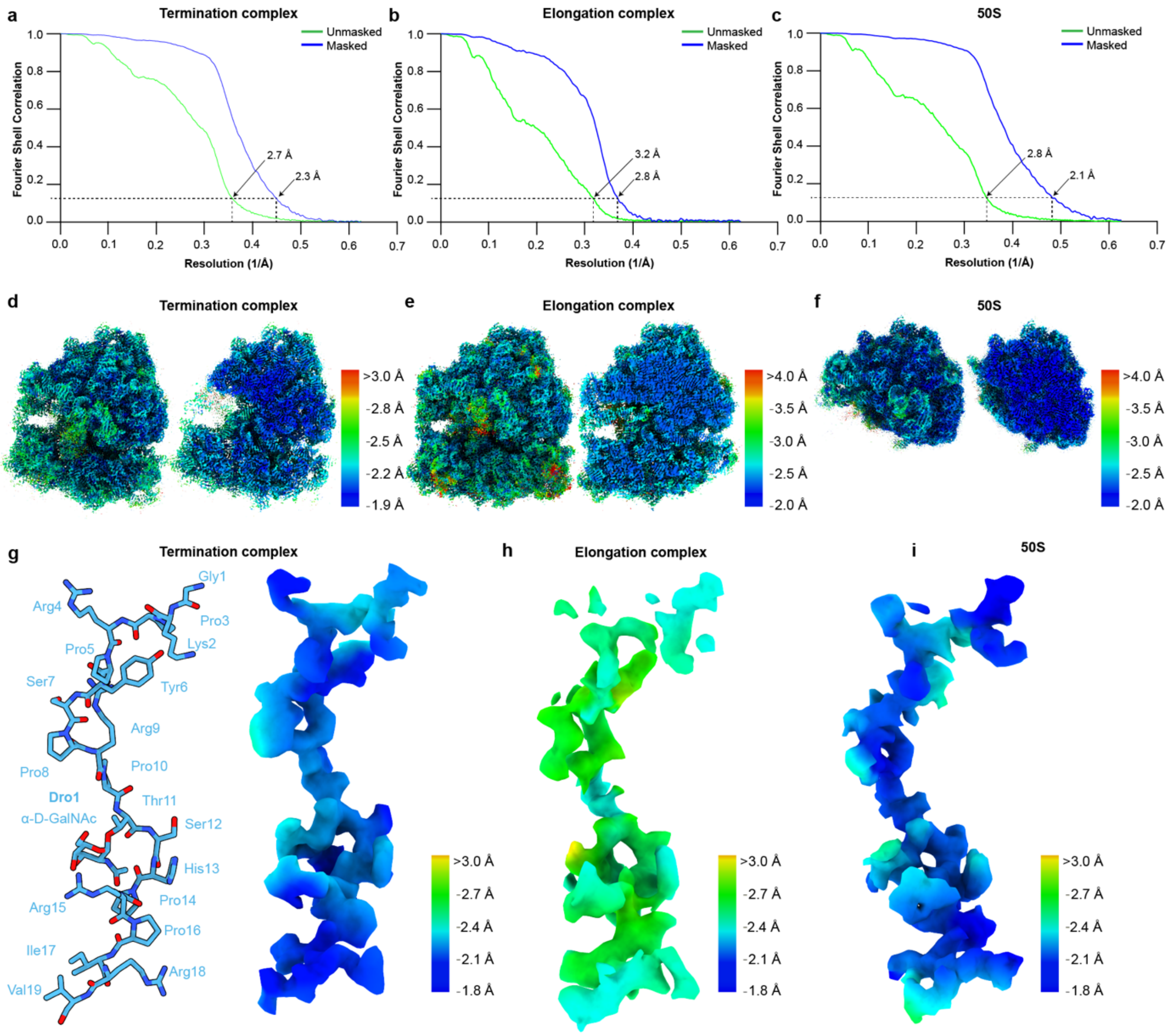


**Supplementary Fig. 3: Cryo-EM data processing. a-c**, Fourier shell correlation (FSC) curve of the (**a**) termination, (**b**) elongation and (**c**) 50S complexes, with unmasked (green) and masked (blue) FSC curves plotted against the resolution (1/Å). **d-f**, Cryo-EM density colored according to local resolution and transverse section for the (**d**) termination, (**e**) elongation and (**f**) 50S complexes. **g-i**, Molecular model of Dro1 (light blue) and corresponding cryo-EM density colored according to local resolution for the (**g**) termination, (**h**) elongation and (**i**) 50S complex.

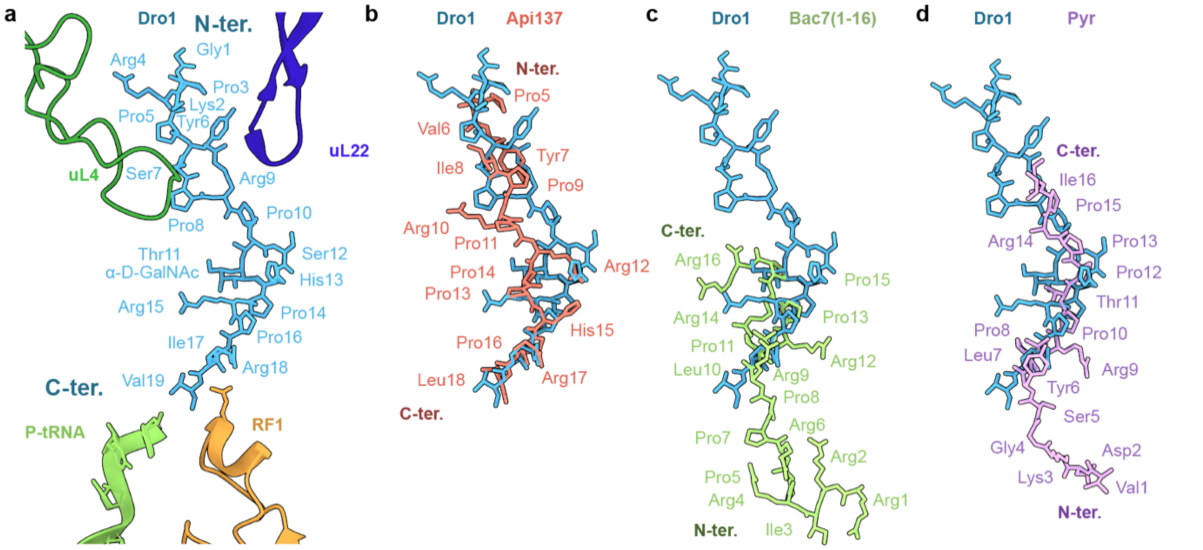


**Supplementary Fig. 4: Dro1 binds with same orientation as Api137, which is distinct from Bac7 and Pyr. a**, Relative position of Dro1 (light blue) compared to P-tRNA (lime), RF1 (orange), uL4 (green) and uL22 (dark blue) within the Dro1-bound termination complex. **b-d**, Superimposition of Dro1 (light blue) from (**a**) with (**b**) Api137 (light red) from the Api137-ArfB complex (PDB ID 6YSS) (Chan et al., 2020), (**c**) Bac7(1-16) (lime) from the Bac7-70S complex (PDB ID 5F8K) (Seefeldt et al., 2016) and (**d**) Pyrrhocoricin (Pyr, purple) from the Pyr-70S complex (PDB ID 5FDV) (Seefeldt et al., 2016).

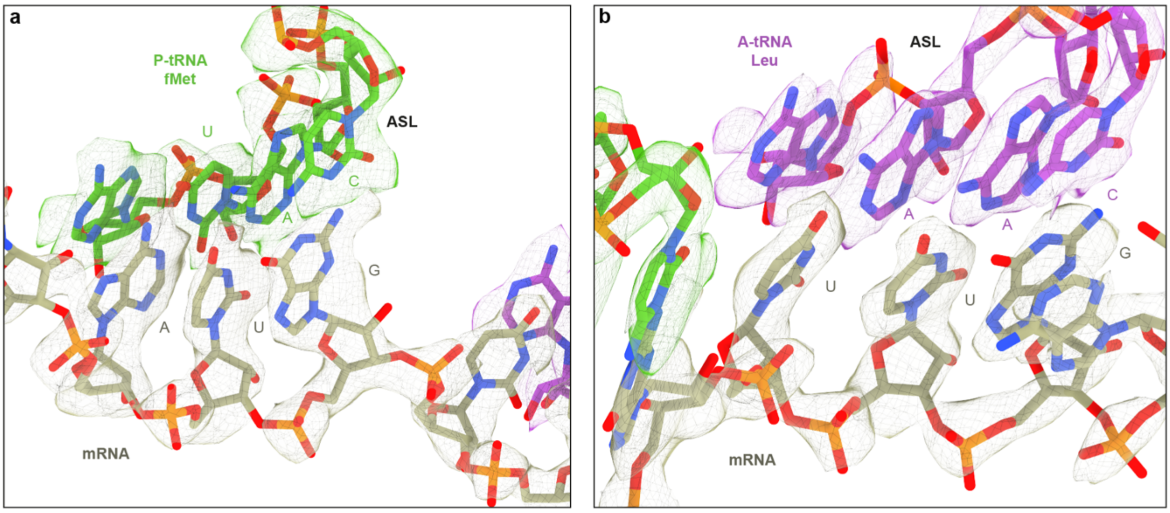


**Supplementary Fig. 5: Anticodon stem loop interaction of the elongation complex. a**, P-site fMet-tRNA UAC anticodon stem loop (lime) interacting with the AUG start codon of the mRNA (light brown) shown with isolated density (mesh). **b**, A-site Leu-tRNA AAC anticodon stem loop (purple) interacting with the UUG codon of the mRNA (light brown) shown with isolated density (mesh).

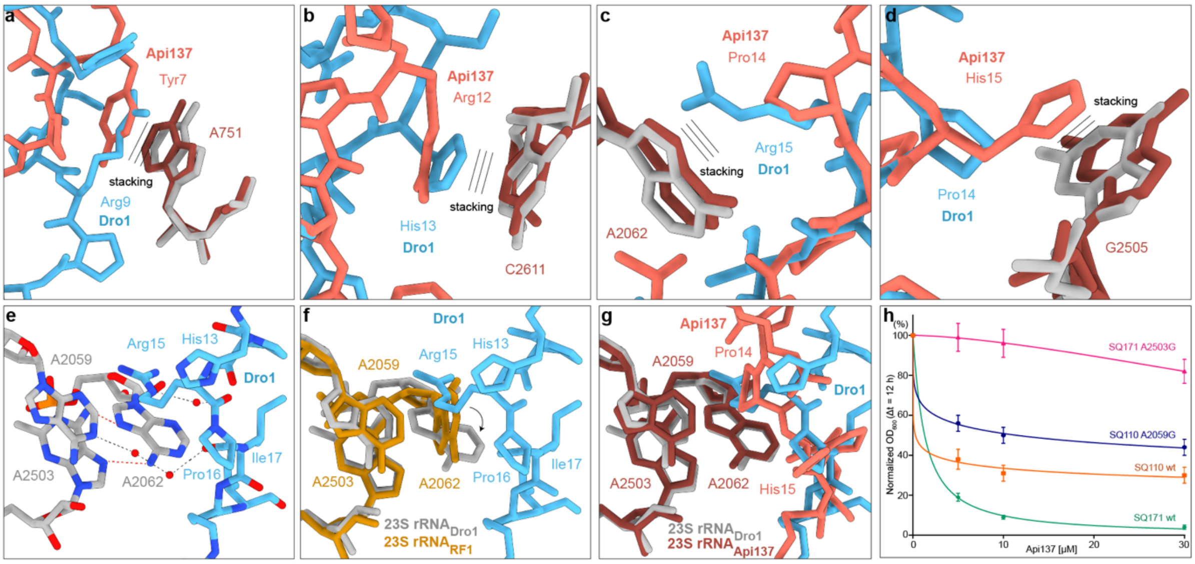


**Supplementary Fig. 6: Api137 interactions compared to Dro1. a-d**, Stacking interactions (indicated as three lines) of sidechains of Api137 (light red, PDB ID 5O2R) (Florin et al., 2017) with 23S rRNA nucleotides (dark red) compared to Dro1 (light blue) stacking interactions with 23S rRNA nucleotides (grey). **e**, Water-mediated interactions of Dro1 (light blue) with surrounding 23S rRNA nucleotides (grey). (**f-g**) Comparison of Dro1 from (**e**) with (**f**) a canonical termination complex (PDB ID 4V63) (Laurberg et al., 2008) and (**g**) Api137 (light red) and corresponding 23S rRNA nucleotides (dark red, PDB ID 5O2R) (Florin et al., 2017). **h**, *in vivo* inhibitory activity of 5 µM, 10 µM and 30 µM Api137 on the growth of *E. coli* SQ110 wt (orange), *E. coli* SQ110 A2059G (blue), *E. coli* SQ171 wt (green) and *E. coli* SQ171 A2503G (pink) in LB medium. For each concentration, inhibition values are the $OD_{600}$ at t = 12 h of the treated culture normalized to the untreated one, considered as 100 %. Error bars represent the standard deviation for three biological replicas and the measurement error of the plate reader. The curves were calculated and plotted by non-linear regression.

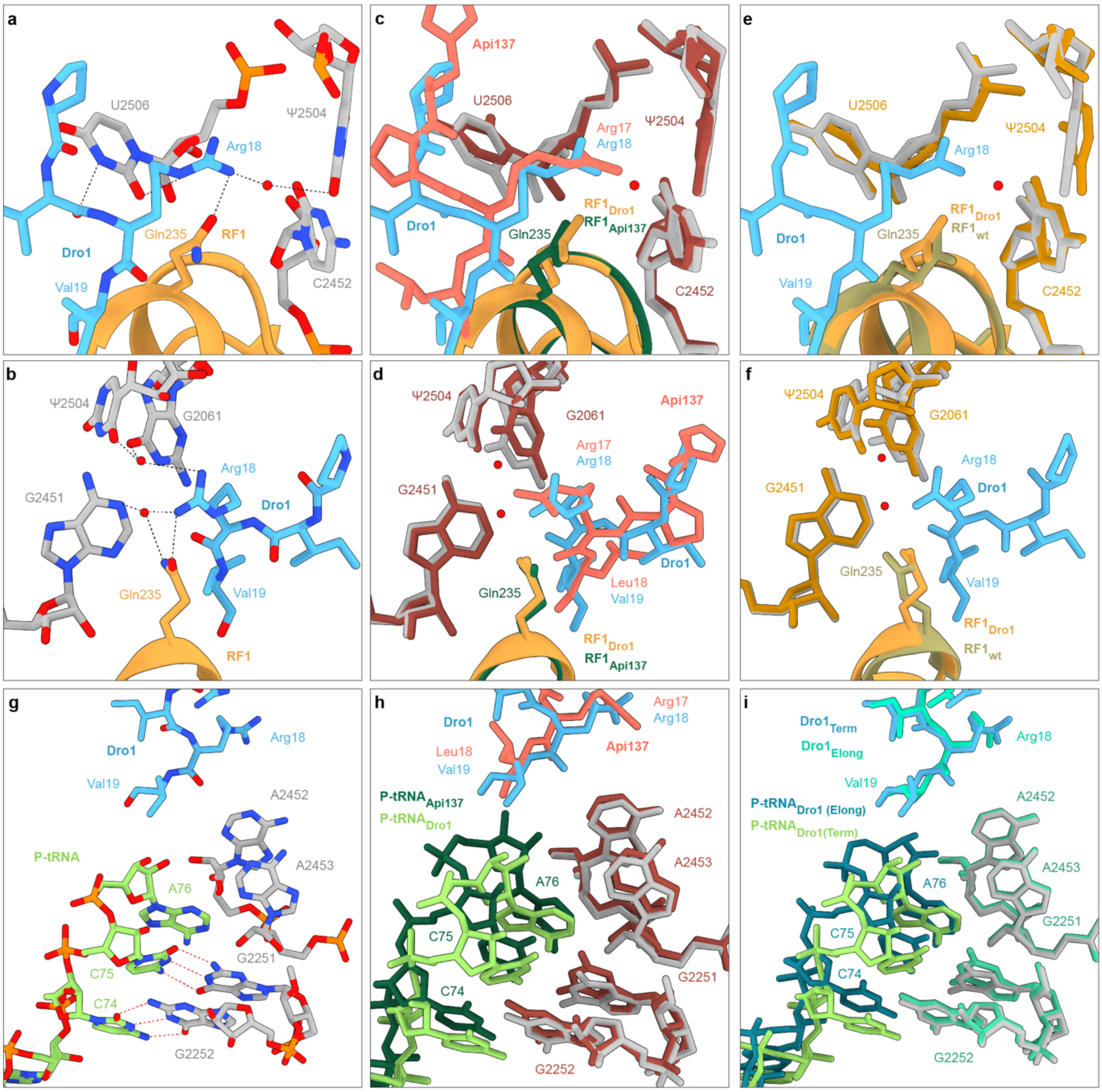


**Supplementary Fig. 7: Interaction of RF1 and P-tRNA with Dro1. a-b**, Water-mediated and direct hydrogen bond interactions of Arg18 of Dro1 (light blue) with surrounding 23S rRNA nucleotides (grey) and Gln235 of RF1 (orange) from two views. **c-f**, Comparison of (**a-b**) with (**c-d**) Api137 (light red, PDB ID 5O2R) (Florin et al., 2017) with RF1 (dark green) and 23S rRNA (dark red) and with (**e-f**) a canonical termination complex RF1 (olive, PDB ID 4V63) (Laurberg et al., 2008) and surrounding 23S rRNA nucleotides (yellow). **g**, CCA-end of a deacylated tRNA (lime) in the P-site in presence of Dro1 is shifted while still establishing base-pairing interactions with G2251 and G2252. **h-i**, Comparison of (**g**) with superimposed deacylated tRNA in the P-site (dark green) and (**h**) the presence of Api137 (light red, PDB ID 5O2R) (Florin et al., 2017) or (**i**) the drosocin-bound elongation complex with deacylated tRNA in the P-site (dark teal) and Dro1 Val19 poorly ordered (teal).

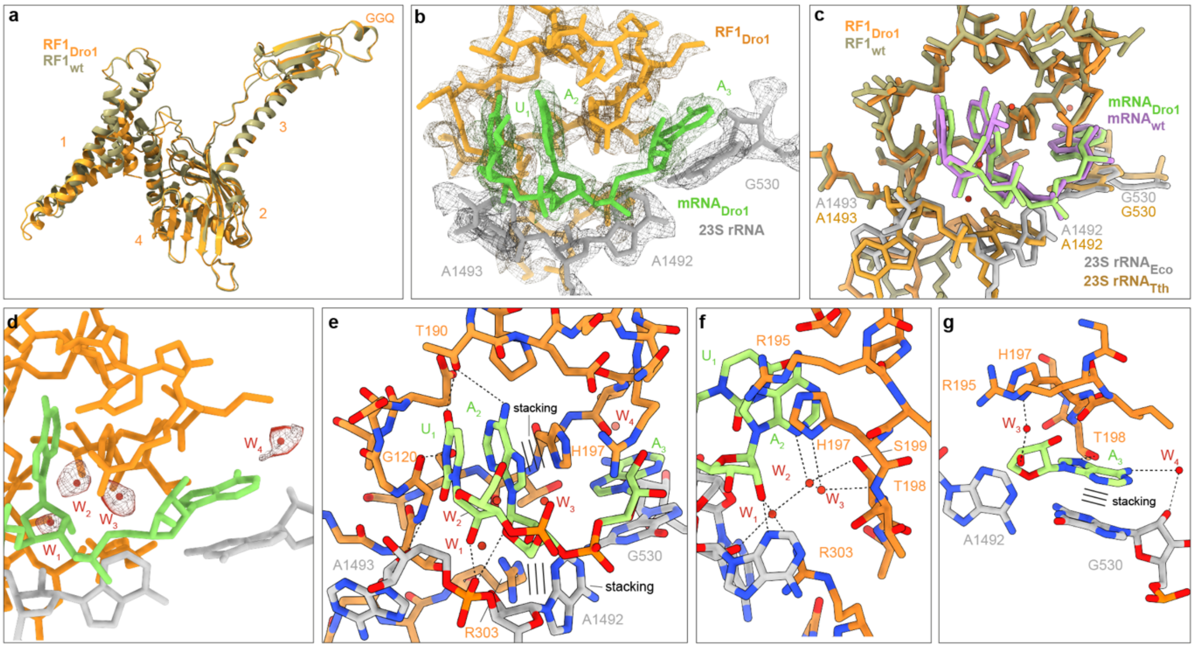


**Supplementary Fig. 8: Stop codon recognition in the drosocin-bound termination complex.** **a**, Superimposition of RF1 in the termination complex (orange) with a canonical RF1 (olive, PDB ID 4V6) (Laurberg et al., 2008). **b**, Isolated density (mesh) for the RF1 (orange), UAA stop codon (lime) of the mRNA and 23S rRNA nucleotides (grey). **c**, Superimposition of RF1 (orange), mRNA (lime) and 23S rRNA nucleotides in the termination complex essential for the stop codon recognition with a canonical termination complex RF1 (olive, PDB ID 4V63) (Laurberg et al., 2008), mRNA (purple) and 23S rRNA nucleotides (grey). **d**, Isolated density assigned to four additional waters molecules in close proximity to RF1 (orange), mRNA (lime), and 23S rRNA nucleotides (grey) within the drosocin-bound termination complex. **e-f**, Potential water-mediated and direct interactions of the UAA nucleotides of the stop codon (lime) with side chains G120, T190, H197, T198, S199 and R303 of RF1 (orange) as well as A1492, A1493 and G530 nucleotides of the 23S rRNA (grey). Hydrogen bonds are indicated by dashed lines and stacking by three lines.

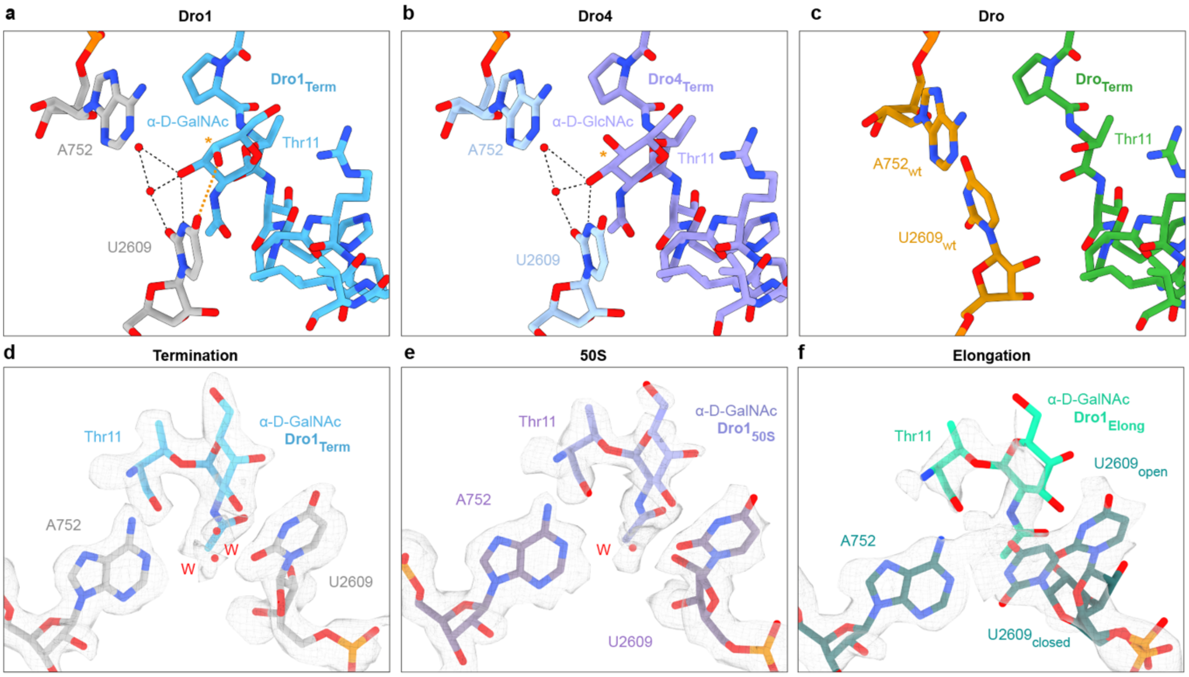


**Supplementary Fig. 9: Sugar modification of drosocin. a-c**, Comparison of (**a**) the α-D-GalNAc modification of Dro1 (light blue) in the termination complex with (**b**) an *in silico* model for the α-D-GlcNAc modification of Dro4 (**b**, light purple) indicating potential loss of a weak hydrogen bond (orange) of C4 hydroxy group of the sugar with C4 hydroxy group of U2609 (grey) as well as (**c**) comparison with an *in silico* model for the unmodified Dro that lacks the Thr-11 glycosylation (green). **d-f**, Isolated density (mesh) for Thr-11 with sugar modification and 23S rRNA nucleotides A752 and U2609 in an open conformation for the (**d**) termination (grey) and (**e**) 50S complexes and (**f**) both open and closed conformations for the elongation complex (dark green).

### Supplementary References


Chan, K.H., Petrychenko, V., Mueller, C., Maracci, C., Holtkamp, W., Wilson, D.N., Fischer, N., and Rodnina, M.V. (2020). Mechanism of ribosome rescue by alternative ribosome-rescue factor B. Nat Commun *11*, 4106.

Florin, T., Maracci, C., Graf, M., Karki, P., Klepacki, D., Berninghausen, O., Beckmann, R., Vazquez-Laslop, N., Wilson, D.N., Rodnina, M.V., *et al.* (2017). An antimicrobial peptide that inhibits translation by trapping release factors on the ribosome. Nat Struct Mol Biol *24*, 752-757.

Laurberg, M., Asahara, H., Korostelev, A., Zhu, J., Trakhanov, S., and Noller, H.F. (2008). Structural basis for translation termination on the 70S ribosome. Nature *454*, 852-857.

Seefeldt, A.C., Graf, M., Perebaskine, N., Nguyen, F., Arenz, S., Mardirossian, M., Scocchi, M., Wilson, D.N., and Innis, C.A. (2016). Structure of the mammalian antimicrobial peptide Bac7(1-16) bound within the exit tunnel of a bacterial ribosome. Nucleic acids research *44*, 2429-2438.
